# Benchmarking Differential Abundance Analysis Methods for Correlated Microbiome Sequencing Data

**DOI:** 10.1101/2022.07.22.501190

**Authors:** Lu Yang, Jun Chen

## Abstract

Differential abundance analysis (DAA) is one central statistical task in microbiome data analysis. A robust and powerful DAA tool can help identify highly confident microbial candidates for further biological validation. Current microbiome studies frequently generate correlated samples from different microbiome sampling schemes such as spatial and temporal sampling. In the past decade, a number of DAA tools for correlated microbiome data (DAA-c) have been proposed. Disturbingly, different DAA-c tools could sometimes produce quite discordant results. To recommend the best practice to the field, we performed the first comprehensive evaluation of existing DAA-c tools using real data-based simulations. Overall, the linear model-based methods LinDA, MaAsLin2, and LDM are more robust than methods based on generalized linear models. The LinDA method is the only method that maintains reasonable performance in the presence of strong compositional effects.

## Introduction

With nearly two-decade research efforts, the human microbiome, the collection of microorganisms and their genetic contents associated with the human body, has been revealed to play a significant role in human health and disease^1^. The human microbiome constantly interacts with environmental and host factors and dynamically evolves over time^2^. It is of great interest to study how the microbiome changes over time and its association with factors such as demographic and lifestyle characteristics, medical history, disease treatment, and various clinical outcomes. To answer these questions, longitudinal designs have been increasingly employed in microbiome studies^3,4^. Compared to case-control and cross-sectional microbiome studies, longitudinal studies, which repeatedly sample the microbiome over a course of time, provide unique opportunities to investigate the dynamics of the microbiome, decipher the species interaction network, and establish a potential causal relationship if the microbiome change precedes the phenotypic change^2^. Statistically, longitudinal studies enjoy higher statistical power and less confounding by using the baseline measurement as the control. Exemplary longitudinal microbiome studies are those from the Integrative Human Microbiome Project (iHMP)^5^, the second phase of the Human Microbiome Project (HMP). iHMP focuses on generating integrated longitudinal datasets and understanding how the microbiome impacts the disease course through a longitudinal view. Besides the longitudinal design, spatial and replicate sampling designs have also been frequently used in microbiome studies^2,4,6^. All these studies generate correlated microbiome data, where the microbiome composition profile derived from the same subject is more similar to each other than those derived from different subjects. Addressing these inherent correlations in microbiome data analysis is critical in obtaining robust and reproducible results.

One central statistical task for microbiome data analysis is differential abundance analysis (DAA), where the goal is to identify the microbial features whose abundance covaries with a variable of interest. The identified microbial features could help improve our understanding of disease mechanisms and be potentially used as biomarkers for disease prevention, diagnosis, prognosis, and treatment selection^7^. With the help of next-generation sequencing technologies, microbiome samples are now routinely profiled by either 16S rRNA gene-targeted sequencing or whole-genome shotgun sequencing^8^. After bioinformatics processing of the sequencing reads, microbiome data can be summarized into a count table, which records the frequencies of the detected microbial features. Depending on the specific pipeline used, these microbial features could be operational taxonomic units (OTUs), amplicon sequence variants (ASVs)^9^, or taxa at different taxonomic ranks. DAA is then performed on the count table together with the metadata describing the sample conditions. DAA of microbiome data raises several statistical challenges including properly modeling the zero-inflated highly skewed abundance distribution^10–12^, addressing the inherent compositional effects^13–15^, and effectively utilizing the phylogenetic relatedness among microbial features^16,17^. In addition to addressing these basic characteristics of microbiome compositional data, DAA of correlated microbiome data also faces the challenge of properly accounting for the correlation structures of non-normally distributed abundance data. Ignoring the correlations could reduce the efficiency of the analysis (analogy to using a two-sample t-test to paired data) or more seriously, produce overly confident results due to exaggeration of the true sample size.

Compared to a plethora of DAA methods developed for independent microbiome data, methods for correlated microbiome data (we label them as “DAA-c”) are relatively under-developed. Nevertheless, in the past decade, several statistical methods were proposed and applied in analyzing correlated microbiome data. These methods could be roughly divided into three categories. **The first category** of methods involves data transformation so that the transformed abundance data are more amenable to modeling by existing statistical methods. Commonly used transformation include log, centered log ratio (CLR)^18,19^, square-root and arcsin-square root transformation^20^. Based on the transformed data, the standard linear mixed-effects model (LMM) is then directly applied^21–23^. The MaAsLin2^24^ package uses LMM as the default method to analyze correlated microbiome data, with additional preprocessing steps (e.g., zero replacement), and several options for normalization and transformation. However, the default total sum scaling (TSS) normalization used in MaAsLin2, could lead to severely inflated type I error under certain scenarios due to strong compositional effects^25^. For example, the increase in the abundance of one dominant microbial species will lead to apparent decreases in the relative abundance of all other species. To address the compositional effects, Zhou et al.^26^ proposed the LinDA method, which corrects the compositional bias after applying LMM on CLR transformed data. Although LMM is computationally efficient and highly interpretable, its normality assumption may not be met for real data due to the severe zero inflation^10–12^. It is unknown whether such assumption violation will lead to reduced power and/or increased type I error. To remedy the drawback of LMM, zero-inflated Gaussian mixed models (ZIGMM)^27^ was proposed. ZIGMM assumes a zero-inflated Gaussian distribution for the transformed abundance data and models the zero and nonzero parts using logistic and linear mixed-effects models, respectively. The LDM method^28^, another linear model-based method based on transformed abundance data, uses permutation to assess the significance so that it is more robust to model misspecification. Different permutation schemes are used in LDM to account for the correlation structure in the data. **The second category** of methods models the TSS-normalized data or proportions using probabilistic distributions with support on [0, 1]. The beta distribution is a popular choice due to its ability to model a wide range of skewed abundance distributions through its two shape parameters. One representative in this category is the two-part zero-inflated beta mixed model (ZIBR)^29^, where the logistic mixed-effects model and the mixed-effect beta regression model are used to model the (structural) zero and non-zero parts, similar in spirit to ZIGMM. **The third category** of methods models the count data through generalized linear mixed-effects models (GLMM). These GLMM methods naturally address the sampling variability of the read counts and use more information than the methods from the first two categories. Thus, they are expected to be more powerful for small-sample studies. However, the major challenges are the computational complexity due to the involvement of integration in the likelihood function and strong model assumptions of the count distribution. The simplest count model is the Binomial or Poisson model. However, real microbiome data exhibit more variability than what is expected by a Binomial or Poisson model. The negative binomial (NB) model, on the other hand, has an extra overdispersion parameter and is more flexible. NB models have been widely used in differential abundance analysis of microbiome data^30,31^. To account for the correlation structure, the mixed-effects negative binomial regression model, which has been implemented in the famous R lme4 package (“glmer.nb” function)^32^, has been applied in practice^33–35^. As an alternative to “glmer.nb”, Zhang, et al. ^36^ developed a flexible and efficient Iterative Weighted Least Squares algorithm to fit the mixed-effects negative binomial regression model (“negative binomial mixed model”, NBMM). Later, Zhang & Yi^37^ extended the NBMM to account for zero inflation and proposed the zero-inflated negative binomial mixed model (FZINBMM). Besides these specialized methods, the zero-inflated negative binomial mixed model can also be fit using the R package GLMMadaptive^38^ and glmmTMB^39^. GLMMadaptive and glmmTMB use an adaptive Gaussian quadrature and Laplace approximation to evaluate the likelihood function, respectively. In addition to the negative binomial model, GLMM with a quasi-Poisson family has also been used for analyzing over-dispersed longitudinal microbiome data. For example, the studies^40,41^ used the Penalized Quasi-Likelihood (GLMMPQL) approach to fit the GLMM with a quasi-Poisson family.

The rising popularity of longitudinal microbiome studies and the availability of multiple DAA-c methods calls for a comprehensive evaluation to provide recommendations and guidance to end-users and tool developers. In contrast to several benchmarking studies of DAA methods^42,43^ for independent microbiome data, no benchmarking studies, to our best knowledge, have been conducted for DAA-c methods for correlated microbiome data. In this study, we propose a real data-based semiparametric simulation framework to perform comprehensive evaluation under diverse biologically relevant settings. We evaluate the performance of DAA-c methods under three commonly seen study designs. These designs include (1) replicate sampling, where each microbiome sample is subject to multiple measurements to reduce noises^44^, (2) matched-pair design, where the microbiome is sampled before and after treatment for each subject^28,45^, and (3) general longitudinal sampling, where the microbiomes of two groups of subjects are sampled at multiple time points^46^. We focus the evaluation on the ability of the DAA-c method to control for false positives and the power to detect true association signals after false discovery rate (FDR) control^47^. The results of the benchmarking study will inform the users to select the most robust method for their studies.

## Results

### The semiparametric simulation approach captures the characteristics of correlated microbiome data

To provide an objective evaluation of DAA-c methods, real microbiome datasets with known truth are the best candidates. However, such datasets are difficult to obtain and even if they do exist, they may only cover limited biologically relevant settings. Therefore, we use simulations, where the ground truth is known, to evaluate the performance of DAA-c methods. To simulate realistic microbiome data, we employ a semiparametric approach, where the baseline compositions are sampled from a reference set of real microbiome data and covariate and confounder effects are then added parametrically (Methods and Fig. S1). This approach circumvents the difficulty in modeling the complex abundance distribution of real microbiome data using statistical models. Previously, we introduced a semiparametric simulation framework for independent data ^48^, where we demonstrated that it could capture the basic characteristics of the real microbiome data such as the sparsity level, mean and variance, and taxon-taxon correlations. In this study, we extended the framework by incorporating within-subject correlations. This is achieved by replicating the subject-level abundance profile *J* times, where *J* is the number of replicates for each subject, followed by adding sample-specific random errors, whose variance (the parameter 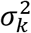, see Supplementary Table 1) controls the correlation strength. Based on the principal coordinate plot (Bray-Curtis distance) on the simulated data, we see that 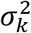 controls the subject-level clustering pattern for the longitudinal microbiome data. As we increase 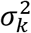 from 1 to 4, the samples from the same subject are less clustered (Fig. S2a). In addition, the approach allows including random slopes so that each subject has its own temporal trajectory (Fig. S2b). We will use this semiparametric framework to evaluate the performance of DAA-c methods for correlated data under diverse settings (Methods, Table 1).

**Table 1.** Simulation settings used in evaluation of DAA-c methods

| Design |  | Setting | Effect size |  |  |  | Between-subject | Within-subject |
| --- | --- | --- | --- | --- | --- | --- | --- | --- |
| | | | $X_i(M_g^X)$ | | $T_i(M_g^T)$ | | variation($S_b^T$) | correlation($\sigma_k^2$) |
| Global null |  | 1 | 0 | 0 | 0 | 0 | 0 | 1 |
|  |  | 2 | 0 | 0 | 0 | 0 | 0 | 4 |
| Replicate sampling | Balanced | 3 | + | ++ | 0 | 0 | 0 | 1 |
|  | Unbalanced | 4 | + | ++ | 0 | 0 | 0 | 1 |
|  | Balanced | 5 | + | ++ | 0 | 0 | 0 | 4 |
|  | Unbalanced | 6 | + | ++ | 0 | 0 | 0 | 4 |
| Matched-pair | Balanced | 7 | 0 | 0 | + | ++ | 0 | 1 |
|  | Unbalanced | 8 | 0 | 0 | + | ++ | 0 | 1 |
|  | Balanced | 9 | 0 | 0 | + | ++ | 0 | 4 |
|  | Unbalanced | 10 | 0 | 0 | + | ++ | 0 | 4 |
| Longitudinal | Balanced | 11 | + | ++ | 0.5 | 0.5 | 0.5 | 1 |
|  |  | 12 | 0.5 | 0.5 | + | ++ | 0.5 | 1 |
|  | Unbalanced | 13 | + | ++ | 0.5 | 0.5 | 0.5 | 1 |
|  |  | 14 | 0.5 | 0.5 | + | ++ | 0.5 | 1 |
|  | Balanced | 15 | + | ++ | 0.5 | 0.5 | 0.5 | 4 |
|  |  | 16 | 0.5 | 0.5 | + | ++ | 0.5 | 4 |
|  | Unbalanced | 17 | + | ++ | 0.5 | 0.5 | 0.5 | 4 |
|  |  | 18 | 0.5 | 0.5 | + | ++ | 0.5 | 4 |

**Table 2.** DAA-c methods evaluated in this study

| Method | Handling zeros | Normalization | Model | R package version | Availability |
| --- | --- | --- | --- | --- | --- |
| ZIGMM | Model(Zero-inflation) | GMPR | Zero-inflated Gaussian mixed model | NBZIMM_1.0 | https://github.com/nyiuab/NBZIMM |
| NBMM | Not necessary |  | Negative binomial mixed models |  |  |
| FZINBMM | Model(Zero-inflation) |  | Zero-inflated negative binomial mixed model |  |  |
| GLMMadaptive | Model(Zero-inflation) |  | Zero-Inflated Poisson/negative binomial mixed model | GLMMadaptive_0.8-0 | https://cran.r-project.org/web/packages/GLMMadaptive/index. |
| glmmTMB | Model(Zero-inflation) |  | zero-inflated negative binomial mixed model | glmmTMB_1.0.2.1 | https://cran.r-project.org/web/packages/glmmTMB/index.html |
| glmer.nb | Model(Zero-inflation) |  | Negative binomial/Possion mixed models | lme4_1.1-26 | https://cran.r-project.org/web/packages/lme4/index.html |
| GLMMPQL | Model(Overdispersion) |  | GLM quasi-Poisson model | MASS_7.3-53 | https://cran.r-project.org/web/packages/MASS/index.html |
| LDM | Not necessary | TSS | Linear model | LDM_1.0 | https://github.com/yijuanhu/LDM |
| LinDA | Pseudo-count | CLR | Linear model | LinDA_0.1.0 | https://github.com/zhouhj1994/LinDA |
| MaAsLin2 | Pseudo-count | TSS | Log linear model | Maaslin2_1.4.0 | https://github.com/biobakery/Maaslin2 |
| ZIBR | Model(Zero-inflation) | TSS | Two-part zero-inflated beta mixed model | ZIBR_0.1 | https://github.com/chvlyl/ZIBR |

### Performance of DAA-c methods under the global null setting

We first study the global null setting, where there are no differential taxa with respect to the covariate *X*_*i*_ (Table 1 settings 1-2, 100 subjects with 2 replicates for each, 500 taxa). We compare the FDR control of various DAA-c methods at the 5% level (Fig. 1, Fig. S3). In this case, FDR is essentially the probability of making any false claims in multiple testing. For stool data, LDM, LinDA and MaAsLin2, all linear model-based methods, could control the FDR close to the target level across settings. In contrast, NBMM, ZIGMM, ZINBMM and glmmadaptive show moderate FDR inflation when the within-subject correlation is low (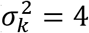), while ZIBR has moderate FDR inflation when the within-subject correlation is high (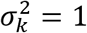). glmernb, GLMMPQL and glmmTMB, on the other hand, have moderate FDR inflation in both settings. For vaginal data, FDR control becomes more challenging. Only LDM and MaAsLin2 can control the FDR to the target level across settings while LinDA shows moderate FDR inflation regardless of the within-subject correlation strength. ZIGMM can control FDR within 10% only when the within-subject correlation is low. All other methods fail to control FDR properly.

**Fig. 1.**
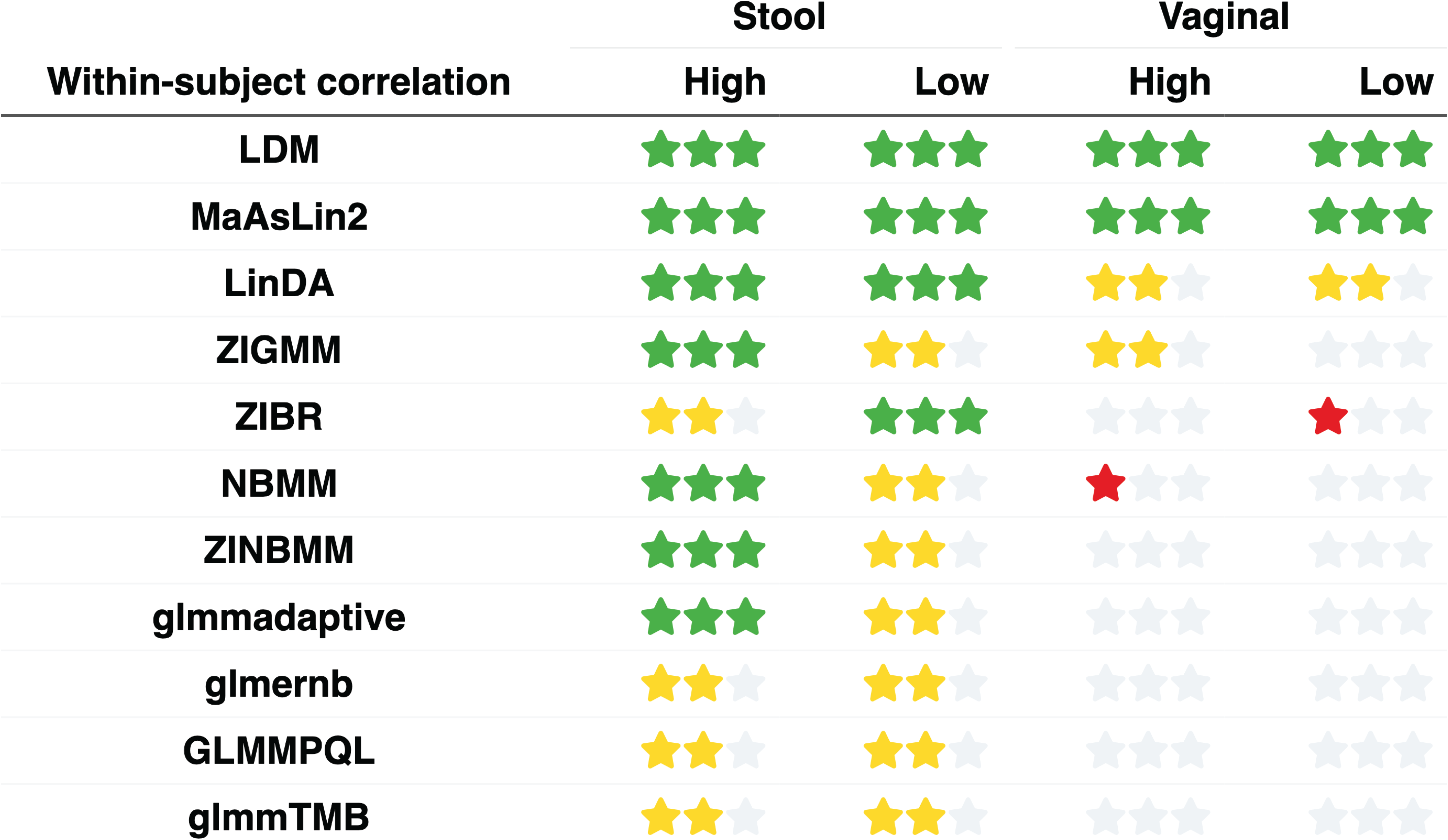
Performance of DAA-c methods under the global null setting for stool and vaginal microbiome data (replicate sampling). Performance is assessed by the observed false discovery rate (FDR) calculated as the percentage of the 1,000 simulation runs making any false discoveries. 3 green stars and 2 yellow stars, 1 red star, and 0 star (all gray) indicate the observed FDR level in [0, 0.05], (0.05,0.1], (0.1,0.2], and (0.2,1], respectively.

### Performance of DAA-c methods under balanced changes

Our next study focuses on the performance of DAA-c methods when the abundance of 10% randomly selected taxa covaries with the treatment covariate (*X*) or the time covariate (*T*) (total sample size: 200, taxa number: 500). In this set of simulations, we simulate balanced changes, i.e., the abundance of those differential taxa increases or decreases in one group randomly. We will test for the effect of *X* for replicate sampling data, the effect of *T* for matched-pair data, and both *X* and *T* for general longitudinal data.

*Performance on replicate sampling data (100 subjects with 2 replicates each, settings 3 and 5, Fig. 2).* For the stool data (Fig. 2a), only LinDA, MaAsLin2 and LDM can control the FDR within 10% across settings. However, LinDA and MaAsLin2 are substantially more powerful than LDM, especially when the within-subject correlation is low. ZIGMM, glmmadaptive, ZINBMM and NBMM can control the FDR within 10% when the within-subject correlation is high and the power is comparable to LinDA and MaAsLin2. But they cannot control the FDR properly when the within-subject correlation is low. ZIBR shows the opposite trend as in the global null setting. In contrast, glmmTMB, GLMMPQL and glmernb show high FDR inflation. For the vaginal data (Fig. 2b), the performance for most methods becomes worse. However, MaAsLin2, LinDA and LDM are still able to control FDR within 10% and MaAsLin2 and LinDA are more powerful than LDM. ZIBR performs well in both FDR control and power when the within-subject correlation is low but cannot control FDR when the within-subject correlation is high. All other methods have severe FDR inflation.

**Fig. 2.**
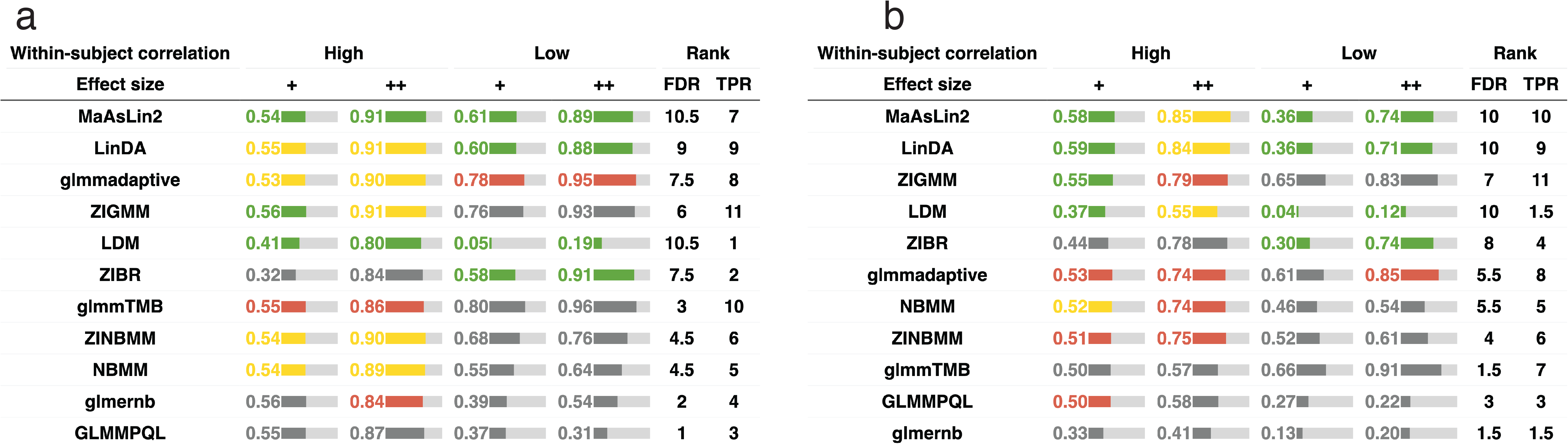
False positive control and power under the replicate sampling design (balanced change setting, **a**: stool and **b**: vaginal). ‘+’, and ‘++’ represent moderate and large effect sizes, respectively. “High” and “Low” within-subject correlations are simulated with 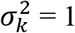 and 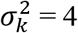, respectively. Green, yellow, red, and gray colors indicate the observed FDR. The green color indicates that the method controls the FDR at the 5% target level (the 95% confidence interval covers 5%). Yellow, red and gray colors indicate the observed FDR level in (0.05-0.1], (0.1, 0.2], and (0.2, 1], respectively. The length of the bar is proportional to the average TPR and the actual value is shown in the bar. FDR and TPR ranks are based on the average FDR and TPR scores across settings. The order of the method is arranged based on the sum of the FDR and TPR ranks.

*Performance on matched-pair data (100 subjects with pre- and post-treatment sample each, settings 7 and 9, Fig. 3)*. Overall, we see a deterioration of the FDR control performance for most methods compared to their performance in the replicate sampling setting (Fig. 3 vs. Fig. 2). For both stool and vaginal data, only LinDA, MaAsLin2 and LDM can control the FDR close to the target level across settings. Again, LinDA and MaAsLin2 are more powerful than LDM. For other methods, only glmmadaptive can control FDR within 10% for the stool data with high within-subject correlation, while other methods fail to control the FDR properly for both stool and vaginal data.

**Fig. 3.**
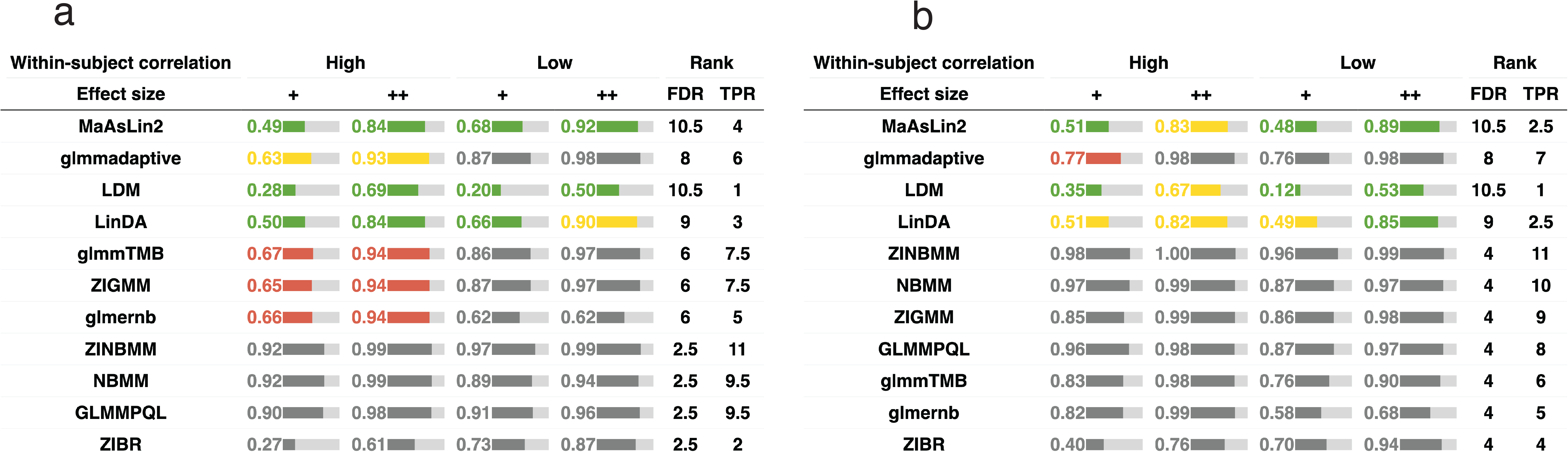
False positive control and power under the matched-pair design (balanced change setting, **a**: stool and **b**: vaginal). ‘+’, and ‘++’ represent moderate and large effect sizes, respectively. “High” and “Low” within-subject correlations are simulated with 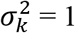 and 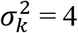, respectively. Green, yellow, red, and gray colors indicate empirical FDR. The green color indicates that the method controls the FDR at the 5% target level (the 95% confidence interval covers 5%). Yellow, red and gray colors indicate the observed FDR level in (0.05-0.1], (0.1, 0.2], and (0.2, 1], respectively. The length of the bar is proportional to the average TPR and the actual TPR is shown in the bar. FDR and TPR ranks are based on the average FDR and TPR scores across settings. The order of the method is arranged based on the sum of the FDR and TPR ranks.

*Performance on longitudinal data (40 subjects each with 5 time points, settings 11-12 and 15-16, Fig. 4)*. We test both the effect of the covariate (*X*) (settings 11-12, Fig. 4ab) and the time variable (*T*) (settings 15-16, Fig. 4cd). Overall, we observe a similar trend as in previous settings. However, there are several noticeable differences. In the case of testing the effect of *X* (Fig. 4ab), for the stool data, LinDA is more powerful than MaAsLin2 when the within-subject correlation is low. For the vaginal data, only MaAsLin2 can control FDR within 10% across settings while LinDA show some FDR inflation (10-20%) when the within-subject correlation is high and the effect size is moderate (“+”). In the case of testing the effect of *T* (Fig. 4cd), for both stool and vaginal data, only LinDA can control FDR to the target level across settings and the FDR control is not at the expense of power. For MaAsLin2, however, we observe some inflated FDR (>10%) when the within-subject correlation is high. In one setting for vaginal data (effect size ‘+’), the FDR inflation of MaAsLin2 is more than 20%.

**Fig. 4.**
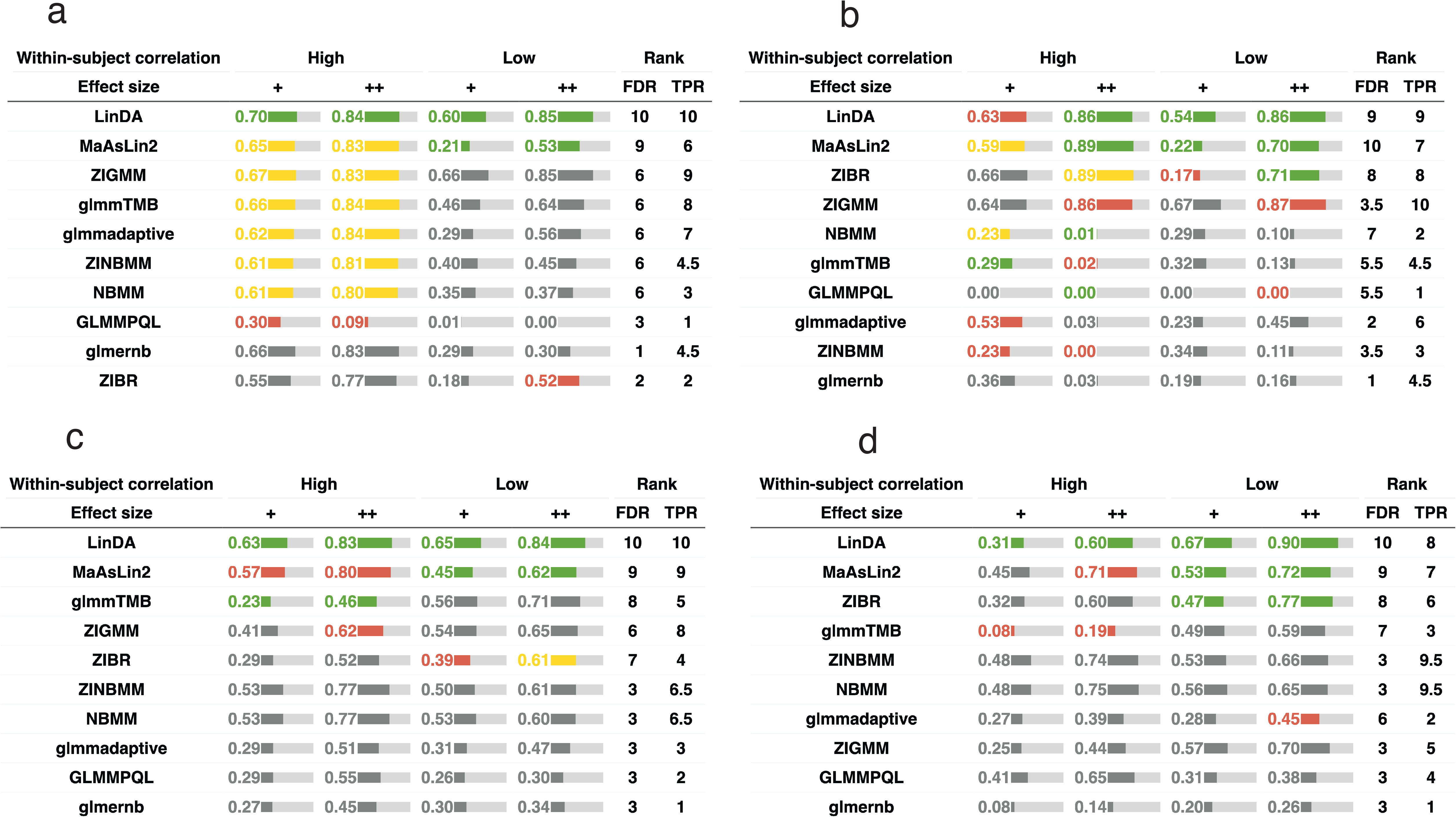
False positive control and power under the general longitudinal design (balanced change setting). **a,b**: testing the group (X) effect [**a**: stool and **b**: vaginal] and **c,d**: testing the time (T) effect [**c**: stool and **d**: vaginal]. ‘+’, and ‘++’ represent moderate and large effect sizes, respectively. “High” and “Low” within-subject correlations are simulated with 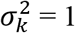 and 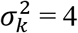, respectively. Green, yellow, red, and gray colors indicate empirical FDR. The green color indicates that the method controls the FDR at the 5% target level (the 95% confidence interval covers 5%). Yellow, red and gray colors indicate the observed FDR level in (0.05-0.1], (0.1, 0.2], and (0.2, 1], respectively. The length of the bar is proportional to the average TPR and the actual TPR is shown in the bar. FDR and TPR ranks are based on the average FDR and TPR scores across settings. The order of the method is arranged based on the sum of the FDR and TPR ranks.

### Performance of DAA-c methods under unbalanced changes

When the differential changes are balanced, the compositional effects are considered to be very moderate. It is interesting to study the performance of the DAA-c methods when the changes are less balanced (i.e., the direction of change is not random) so that the compositional effects are strong. In this new set of simulations, we let the direction of change for those differential taxa be the same. Although such a setting may not be common in practice, it could be used to test the limit of DAA-c methods in addressing compositional effects. We repeat similar analyses under the replicate sampling (Fig. S4, settings 4 and 6), matched-pair (Fig. 5, settings 8 and 10), and longitudinal (Fig. 6, settings 13-14 and 17-18) designs.

**Fig. 5.**
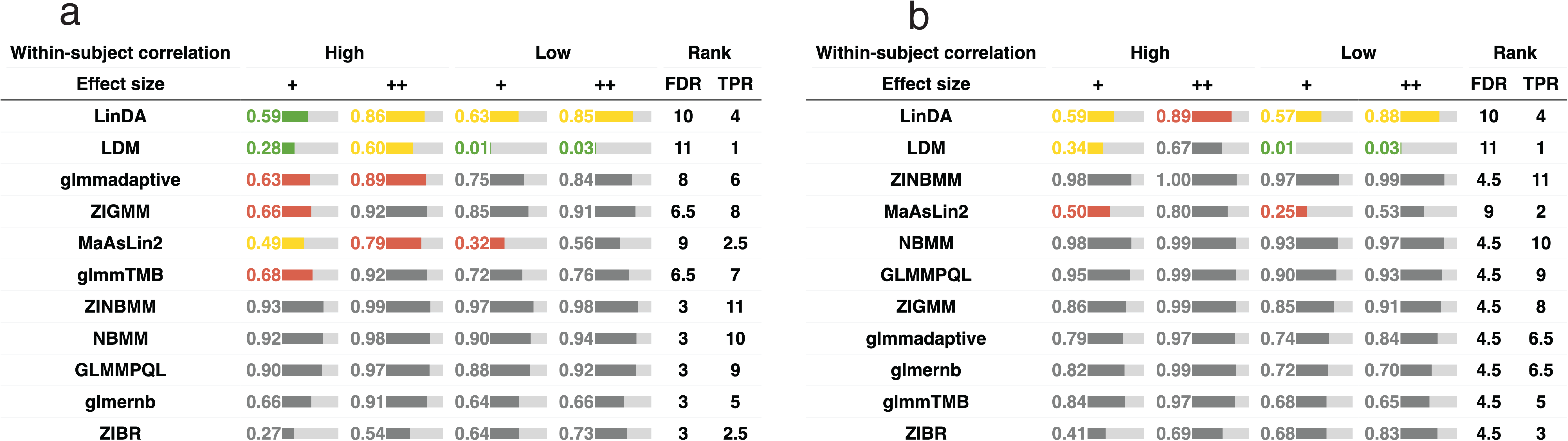
False positive control and power under the matched-pair design (unbalanced change setting, **a**: stool and **b**: vaginal). ‘+’, and ‘++’ represent moderate and large effect sizes, respectively. “High” and “Low” within-subject correlations are simulated with 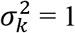 and 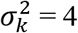, respectively. Green, yellow, red, and gray colors indicate empirical FDR. The green color indicates that the method controls the FDR at the 5% target level (the 95% confidence interval covers 5%). Yellow, red and gray colors indicate the observed FDR level in (0.05-0.1], (0.1, 0.2], and (0.2, 1], respectively. The length of the bar is proportional to the average TPR and the actual TPR is shown in the bar. FDR and TPR ranks are based on the average FDR and TPR scores across settings. The order of the method is arranged based on the sum of the FDR and TPR ranks.

**Fig. 6.**
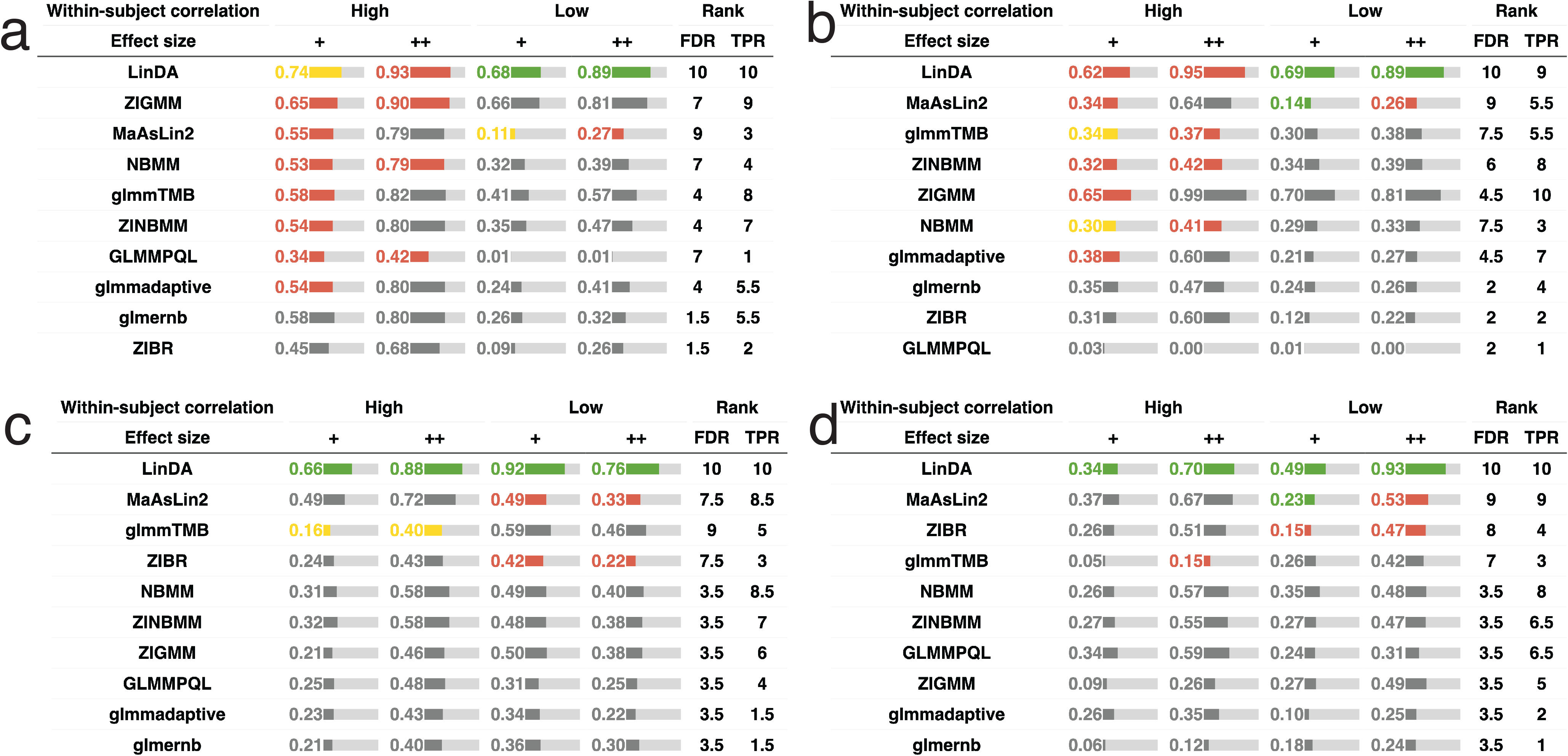
False positive control and power under the general longitudinal design (unbalanced change setting). **a,b**: testing the group (X) [**a**: stool and **b**: vaginal] and **c,d**: testing the time (T) effect [**c**: stool and **d**: vaginal]. ‘+’, and ‘++’ represent moderate and large effect sizes, respectively. “High” and “Low” within-subject correlations are simulated with 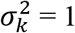 and 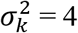, respectively. Green, yellow, red, and gray colors indicate empirical FDR. The green color indicates that the method controls the FDR at the 5% target level (the 95% confidence interval covers 5%). Yellow, red and gray colors indicate the observed FDR level in (0.05-0.1], (0.1, 0.2], and (0.2, 1], respectively. The length of the bar is proportional to the average TPR and the actual TPR is shown in the bar. FDR and TPR ranks are based on the average FDR and TPR scores across settings. The order of the method is arranged based on the sum of the FDR and TPR ranks.

Unsurprisingly, compared to their performance in the balanced change scenario, the FDR control for all DAA-c methods becomes much worse in the unbalanced settings. None of the methods, including LinDA, MaAsLin2 and LDM, could control the FDR within 10% across settings (Fig. S4, Fig. 5-6). The FDR control performance deteriorates with increasing effect size (‘+’ vs. ‘++’), indicating the challenge of differential abundance analysis in the presence of strong compositional effects. The FDR control is poorer for most methods in testing the treatment effect for matched-pair data (Fig. 5) or testing the time effect in longitudinal data (Fig. 6cd). A lower within-subject correlation and a lower microbial diversity (vaginal) also tend to decrease the FDR control performance for many methods.

Among the three best-performing methods in the balanced change settings (LinDA, MaAsLin2 and LDM), LinDA has overall the best FDR control: it can control the FDR within 20% for all settings and within 10% when the within-subject correlation is low. Notably, LinDA can control the FDR to the target level when testing the time effect for the longitudinal data while other methods fail to control FDR properly. The power of LinDA is also comparable to competing methods.

For MaAsLin2, we used the default TSS normalization in comparison. It is interesting to see if its FDR control performance improves with alternative normalization methods. We thus replace the default TSS normalization in MaAsLin2 with Geometric Mean Pairwise Ratios (GMPR)^49^, Trimmed mean of M values (TMM)^50^ and cumulative sum scaling (CSS) normalization^51^. We can see that the FDR control of MaAsLin2 does improve significantly, but it still does not perform as well as LinDA (Fig. S5).

### Effect of the sample size and the number of taxa on the performance of DAA-c methods

In practice, many microbiome studies are conducted with small sample sizes. Moreover, in differential abundance analysis at a higher taxonomic rank, the number of tested taxa may be small. Thus, we want to check how the DAA-c methods perform when the sample size or the number of taxa is small. We use the stool data under the replicate sampling design to study the effect of a small number of samples and taxa.

We first decrease the sample size to 40 (20 subjects with 2 samples each) while the number of taxa to be tested remains at 500. Compared to the results with a sample size of 200 (Fig. 2), we observe a significant decrease in the performance of FDR control for most methods (Fig. S6ab). Those count-based methods perform poorly with both severe FDR inflation and low power. In contrast, those linear model-based methods, MaAsLin2, LinDA and LDM, are more robust to small sample sizes when the changes are more balanced (Fig. S6a). MaAsLin2 and LinDA are substantially more powerful than LDM. MaAsLin2 has the best FDR control performance in this scenario while LinDA has some noted FDR inflation in one setting. These results suggest that simpler models (i.e., linear models) may be preferred over complex models (i.e., count-based generalized linear model) when the sample size is small. When the changes are unbalanced (Fig. S6b), none of the methods including LinDA can control FDR with adequate power when the within-subject correlation is high, and the effect size is large. When the within-subject correlation is low, LinDA excels in both FDR control and power while other methods perform poorly.

We next decrease the number of taxa to 50 by keeping the most abundant taxa in the analysis (Fig. S6cd). Again, most evaluated methods show decreased performance in FDR control, due to the increased compositional effects with a smaller number of taxa. When the changes are balanced (Fig. S6c), MaAsLin2 and LinDA perform much better than other methods with MaAsLin2 being able to control the FDR to the target level across settings. However, when the changes are unbalanced (Fig. S6d), MaAsLin2 has a significantly higher FDR than LinDA.

### Computational efficiency and performance summary

Computationally efficiency is an important factor influencing a user’s choice of methods. Therefore, we compare the computational speeds of the evaluated DAA methods based on the simulated data. We find that only 6 out of 11 methods (ZINBMM, NBMM, ZIGMM, GLMMPQL, MaAsLin2, LinDA) can complete the analysis of a moderate-sized microbiome dataset (100 subjects, 2 replicates each, 500 taxa) within 10 minutes on our computer system (x86_64-pc-linux-gnu (64-bit) Red Hat Enterprise Linux Server 7.9, Intel(R) Xeon(R) CPU E5-2698 v4 @ 2.20GHz, 8GB running memory). LinDA and MaAsLin2 are the fastest methods that can finish the analysis within 1 minute (Fig. S7).

Finally, we summarize the DAA performance using different metrics based on our simulation studies (Fig. 7). For each evaluation metric, we classify each method as ‘good’, ‘intermediate’ or ‘poor’ (see legend in Fig. 7). Although it is difficult to capture the full complexity of the evaluation based on a crude categorization, the heatmap provides a convenient way to convey the major findings in the simulation studies. It is easy to spot that LinDA, LDM and MaAsLin2 have much better FDR control performance than the other methods. LinDA is the only method that can control FDR reasonably well across settings while LDM and MaAsLin2 can have poor FDR control when the compositional effects are strong (e.g., unbalanced change settings). Remarkably, LinDA is overall more powerful than competing methods.

**Fig. 7.**
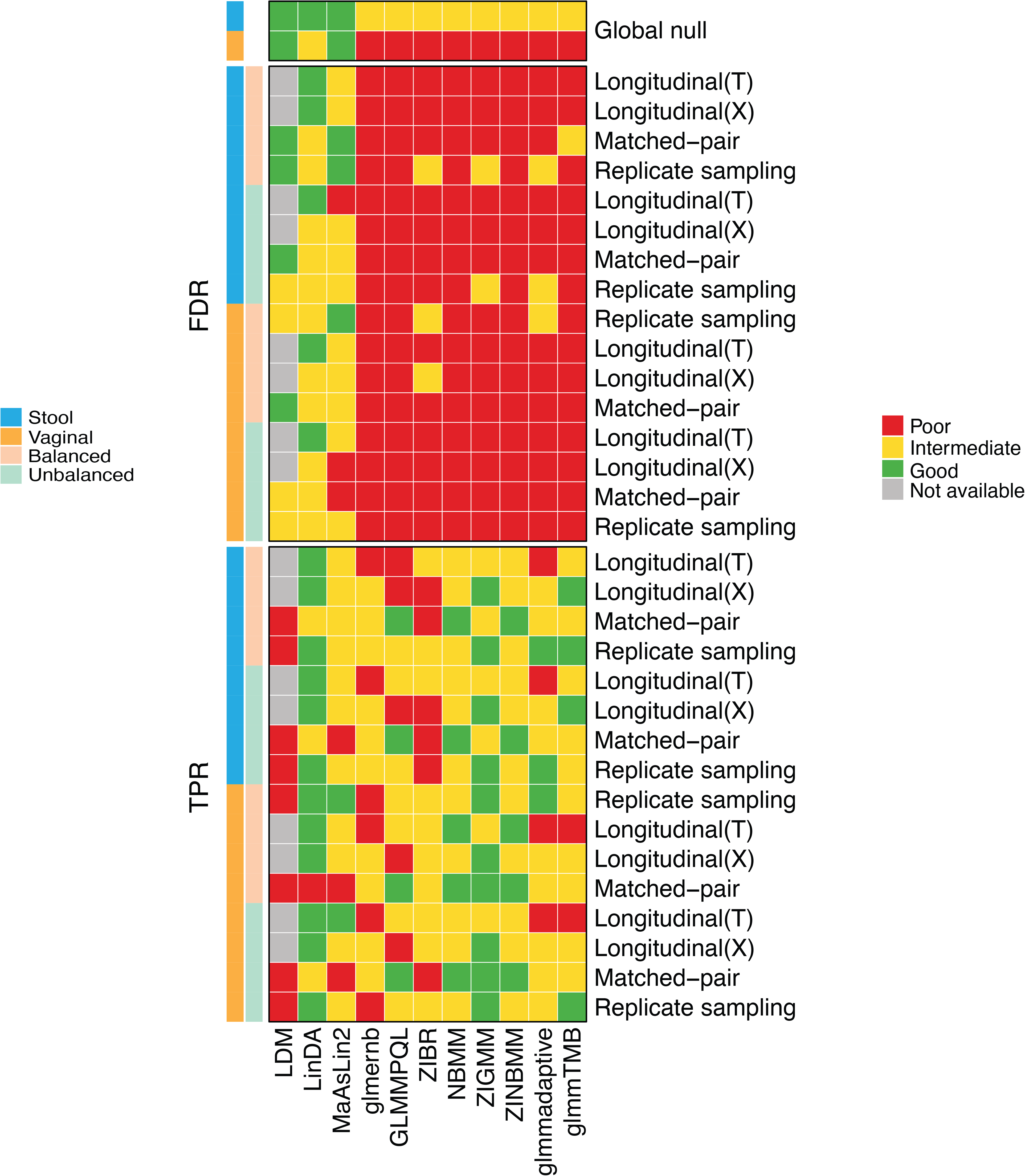
Performance summary of DAA-c methods based on various evaluation metrics. For each metric, the performance is categorized into “Good”, “Intermediate” and “Poor”. For FDR control, “Good”, “Intermediate” and “Poor” represent observed FDR (averaged over two effect sizes and low and high within-subject correlations) in (0,0.05], (0.05,0.2], (0.2,1]. For power, “Good”, “Intermediate” and “Poor” represent top 1/3, mid 1/3, and bottom 1/3 based on their TPR rank. For computational speed, “Good”, “Intermediate” and “Poor” correspond to a computational time (mins/per run) < 1, (1 - 60] and > 60, respectively. LDM currently does not support the general longitudinal design and is colored gray in the corresponding rows in the heatmap.

### Discovery patterns on real correlated microbiome datasets

We next apply the evaluated DAA-c methods to three publicly available datasets^6,52,53^ (“Method”), which are examples of replicating sampling (“Smoker2010”), matched-pair (“Nicholas2013”) and longitudinal (“IBD2017”) designs. Since the ground truth is unknown for the three real datasets, we aim to assess whether the discovery pattern on the real datasets reflects what we have observed in the simulation study. We first evaluate the FDR control of DAA-c methods under the global null by shuffling the sample labels (1,000 times) to disrupt the differential signals. ZIBR currently do not support an unequal number of samples for each subject and LDM does not support general longitudinal designs, so they were excluded from the comparison on the third dataset. For the first dataset, the smoking status of the subjects were permuted. For the second dataset, the pre-cleaning and post-cleaning status labels for each subject were permuted. For the third dataset, group labels were shuffled and time points were permuted within the subjects. Any differential taxa identified from the permuted datasets are considered to be false positives. Therefore, if we use the Benjamini-Hochberg FDR control procedure to identify differential taxa at 5% FDR, we expect to see on average 5% of the permuted datasets to have any false findings. Consistent with the simulation results, most methods do not perform satisfactorily in controlling for false positives (Fig. 8ab). For the “Smoker2010” dataset, LDM, LinDA, MaAsLin2, GLMMPQL, NBMM and ZINBMM can control FDR to the 5% target level (Fig. 8a) and the percentage of differential taxa range from 0% to 12% (median: 0%, Fig. 8b) on the permutated datasets. ZIGMM has slight FDR inflation (8%) with the percentage of differential taxa ranging from 0% to 14% (median 0%). In contrast, all other methods have seriously inflated FDR (>20%), and the percentage of differential taxa ranges from 0% to 20% (median: 3%). For the “Nicholas2013” dataset, LinDA, MaAsLin2 and LDM stand out among their competitors with an observed FDR less than 5% and the percentage of differential taxa ranging from 0% to 6% (median: 0%). All other methods fail to control FDR properly. For the “IBD2017” dataset, only MaAsLin2 and LinDA control the FDR around 5% for both testing the group effect and the group-time interaction effect.

**Fig. 8.**
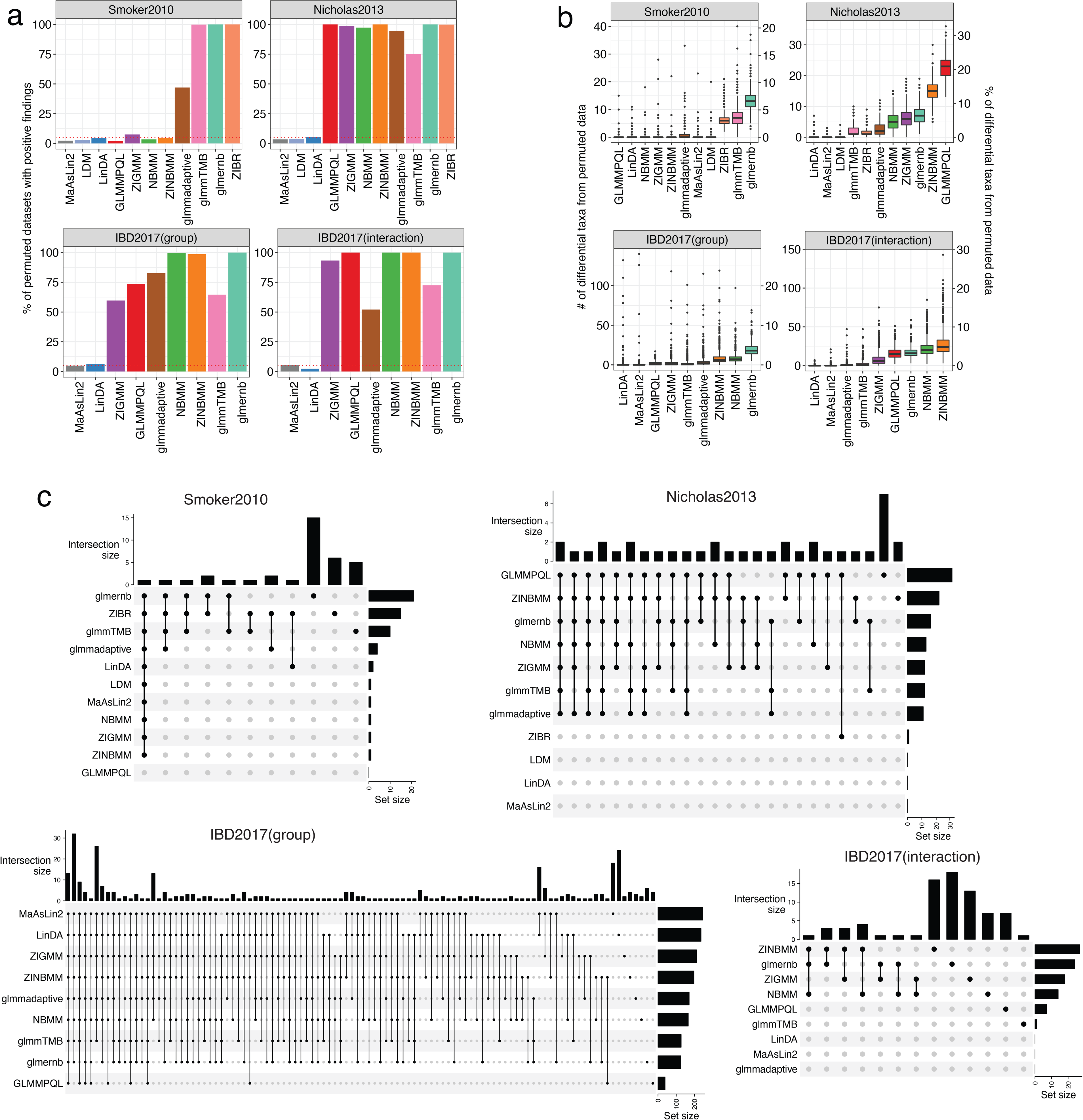
Evaluation of longitudinal differential abundance analysis (DAA-c) methods based on three experimental datasets. **a,b**. Performance evaluation under the global null setting. **a**. Performance is assessed by the observed FDR level calculated as the percentage of the 1,000 simulation runs making any false discoveries. **b**. Boxplot showing the number of differential taxa at 5% FDR (left y-axis) and the percentage of differential taxa (right y-axis) based on 1000 permuted datasets. **c**. Overlaps of significant taxa (5% FDR) between DAA-c methods on the real datasets. Set size means the total number of differential taxa discovered by each method. Intersection size means the number of differential taxa commonly found by the methods indicated by the black dots.

Next, we compare the numbers of identified differential taxa at 5% FDR for those DAA-c methods and study their overlaps based on the original real datasets (Fig. 8c). The results at 10% FDR can be found in Fig. S8. As expected, methods that do not control for false positives tend to find more differential taxa (horizontal bars, Fig. 8c, with one exception), many of which are unique to themselves (right side of the top vertical bars, Fig. 8c). For the “Smoker2010” dataset, LinDA identifies 2 OTUs associated with smoking while the other methods that control for false positives identify 1 or 0 OTU. For the “Nicholas2013” dataset, LDM, LinDA, and MaAsLin2, the three methods that have good false positive control on the permuted datasets, do not identify any significant genera. Although other methods identify a few, the large FDR inflation on the permuted datasets casts doubt on the credibility of the identified genera. For the “IBD2017” dataset, when testing the group effect (ICDr vs. control), MaAsLin2 and LinDA identify more OTUs than the other methods, indicating that their well-controlled FDR may not be at the expense of the power. It is well known that the gut microbiome of IBD patients is very distinguishable from that of healthy controls^54^, supporting the findings of MaAsLin2 and LinDA. However, when testing the interaction between group and time, neither MaAsLin2 nor LinDA identifies any significant interactions. Given the fact that interaction detection usually requires a large sample size, such negative findings are not surprising. Overall, the detection patterns on the real datasets are consistent with those from the simulation study.

## Discussion

Differential abundance analysis is one of the most fundamental statistical tasks in microbiome data analysis. Given the rising popularity of complex study designs involving temporal^55^, spatial^56^, and repeated sampling^4^ of the microbiome, statistical tools that could properly address the correlation structure in microbiome data are much needed. Although there are several tools for differential abundance analysis of correlated microbiome data (DAA-c)^26–29,36–38^, their performance has not been evaluated independently by a large-scale benchmarking study. It is unclear whether these methods can control for false positives while retaining sufficient power for real microbiome datasets under diverse settings. In this study, we thus conducted a large-scale simulation study to objectively evaluate the performance of the major existing DAA-c methods under a wide range of settings. We aim to identify and recommend the most robust DAA-c tool to the field. To achieve this end, we designed a semiparametric simulation framework for realistic microbiome data generation in an extension of our previous work for independent data^48^. The proposed simulation framework circumvents the difficulty in properly modeling the zero-inflated, highly skewed abundance distribution by drawing random samples from a large reference dataset. Covariate and confounder effects are added parametrically to generate correlated microbiome data. We show that the generated data capture the basic characteristics of microbiome data^48^ and thus are more suitable for benchmarking DAA-c methods than parametric model-simulated data.

Based on the proposed simulation framework, we performed the evaluation covering both low- and high-diversity microbial communities. We show that the FDR control performance of evaluated DAA-c methods varies tremendously, and most DAA-c methods are still not satisfactory. In fact, none of the evaluated methods could control the FDR to the target level across all settings. The FDR control is more difficult for the low-diversity community such as the vaginal microbiome since the data are much sparser. The FDR control also deteriorates as the effect size becomes stronger and the direction of change becomes less balanced since these conditions result in stronger compositional effects. For methods that model the count distribution using negative binomial distribution or zero-inflated negative binomial distribution (GLMMPQL, glmmadaptive, NBMM, ZINBMM, glmernb, glmmTMB), they tend to have severe FDR inflation even when the compositional effects are small, indicating that the assumed count model may still not be able to capture the abundance variation adequately^57,58^. Their performance worsens as the sample size becomes smaller probably because the asymptotic distribution of the test statistic, on which the p-value calculation depends, is not accurate for small sample sizes. These methods tend to perform less well when the within-subject correlation is lower, indicating that they probably overfit the correlations. For ZIBR, which models the proportion data using zero-inflated beta distribution, large FDR inflation was observed in most settings and the FDR inflation became more serious when the within-subject correlation was higher, indicating that the zero-inflated beta distribution may not fit the data well, either. In contrast, those methods based on data transformation and linear models (LinDA, MaAsLin2 and LDM) are more robust and they have much better FDR control than the methods that model counts and proportions. When the changes are more balanced, they can control the FDR close to the target level. In terms of power, MaAsLin2 and LinDA are more powerful than LDM especially when the whin-subject correlation is low. However, when the changes are less balanced, the FDR control of MaAsLin2 and LDM deteriorates since they do not specifically address the compositional effects other than using robust normalization factors. In contrast, LinDA, which corrects the bias due to compositional effects, has substantially better FDR control than MaAsLin2 and LDM when the compositional effect is strong. The improved FDR control is not at the excessive expense of the power. However, FDR inflation was still noted for LinDA when the within-subject correlation was high, and the sample size was small.

Based on the evaluation, we find that LinDA has the best trade-off between FDR control and power across settings. Due to potential FDR inflation under some settings, we highly recommend using a standard FDR level such as 5% and resisting the temptation to raise the level to find “signals”. As an alternative to LinDA, MaAsLin2 can also be applied when the compositional effects are moderate. However, some diagnostics for compositional effects are needed if MaAsLin2 is chosen to perform differential abundance analysis. The magnitude of compositional effects can be detected by an effect size plot for those differential taxa. If there are many taxa with the same direction of change, applying MaAsLin2 is not recommended. Otherwise, MaAsLin2 can be used due to its power advantage.

When simulating longitudinal data, we did not include a time-covariate interaction term. However, in real applications, the interaction term may be of major interest. We have implemented the simulation framework in our R GUniFrac package (*SimulateMSeqC* function) and the user can easily modify the function to incorporate an interaction term.

Finally, we comment that there is still plenty of room for novel methodological development for differential abundance analysis of correlated microbiome data. Our simulation framework can be easily re-used to evaluate the performance of new methods.

## Methods

### A semiparametric simulation framework for realistic correlated microbiome data generation

Traditional microbiome data simulators are usually based on parametric models such as Dirichlet-multinomial model^59,60^ and logistic normal multinomial model^61^. The sample space is thus determined by a small set of parameters. Due to the complexity of the microbiome data, existing parametric models may fail to capture the full distributional characteristics of the data. To generate more realistic data, we adopt a semiparametric approach, where we draw random samples from a large reference microbiome dataset (non-parametric part) and add covariate/confounder effects parametrically (parametric part). Basically, for each drawn reference sample, we infer the underlying composition based on an empirical Bayesian model and add covariate/confounder effects to the composition vector via a log linear model. Once the true underlying composition is obtained, sequence reads are generated using a multinomial model. By using the real microbiome data as the template, our method circumvents the difficulty in modeling the complex inter-subject variation of the microbiome composition.

The basic steps of the semiparametric simulation framework are depicted in Fig. S1. Specifically, we use the following steps to generate the longitudinal microbiome data with *I* subjects belonging to two groups and *J* evenly spaced time points for each subject:

1. Build a reference dataset. The reference dataset is a collection of microbiome sequencing samples from a specific body site of a study population. It should be large enough to capture the main compositional variation in the population of interest. Microbiome datasets from those large-scale population-level studies such as Human Microbiome Project (HMP)^5^ and American Gut Project (AGP)^62^ are all good choices. The reference datasets used in the simulation are the human stool and vaginal microbiome datasets from HMP with basic filtering to remove extremely rare taxa (prevalence < 10% or max proportion < 0.2%), resulting in 295 samples and 2094 taxa, and 381 samples and 781 taxa for the stool and vaginal dataset, respectively. The human stool and vaginal microbiome are chosen to represent a high- and low-diversity microbial community, respectively.
2. Sample *a posteriori* the underlying composition of the reference samples based on the observed counts using an empirical Bayes approach.

a. Assume an informative Dirichlet prior for the underlying composition, estimate the Dirichlet hyperparameters (γ_*k*_) based on the observed counts (*C*_*ki*_, 1 ≤ *k* ≤ *M*, 1 ≤ *i* ≤ *N*) using the maximum likelihood estimation (R package “dirmult”). The posterior distribution of the underlying composition for sample *i* is then a Dirichlet distribution with parameter 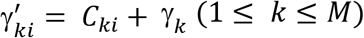.
b. Obtain a posterior sample of the underlying composition for each reference sample based on the posterior Dirichlet distribution. Denote *P*_*ki*_ (1 ≤ *k* ≤ *M*, 1 ≤ *i* ≤ *N*) be the proportion for the *k*th taxon in the *i*th subject.
3. Generate the absolute abundance (*A*_*ki*_) by multiplying a factor *S*_*i*_ representing the microbial load at the sampling site, i.e., *A*_*ki*_ = *P*_*ki*_*S*_*i*_, where log (*S*_*i*_) ∼ *N*(0,1) without loss of generality.
4. Given *I* subjects and *J* time points, randomly draw *I* samples based on the absolute abundance data generated in the last step. Replicate the absolute abundance profile of each subject *J* times. Denote *A*_*kij*_ as the absolute abundance for the *k*th taxon in the *i*th subject at the *j*th time point.
5. Generate the time and group covariates and a confounder. A binary group covariate *X*_*i*_ (1 ≤ *i* ≤ *I*) is created by dichotomizing a latent variable 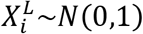 using some cutoff value to achieve the specified group sizes. The confounder is generated by 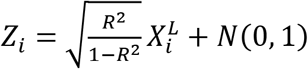, where *R* is the desired correlation between *Z*_*i*_ and 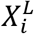. The time covariate *T*_*ij*_ (1 ≤ *i* ≤ *I*, 1 ≤ *j* ≤ *J*) is set as *j* − 1.
6. Given *K* taxa included in the analysis, generate their coefficients for *X*_*i*_, *Z*_*i*_, *T*_*ij*_, respectively, which are 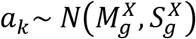, 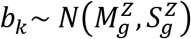, and 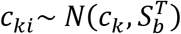, where 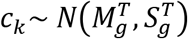, for 1 ≤ *k* ≤ *K*, 1 ≤ *i* ≤ *I*. The interpretation of the parameters 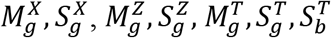 can be found in Supplementary Table 1. Note that the time coefficient *c*_*ki*_ could vary by subject so that each subject could have its own trajectory (random slope). Non-differential taxa are simulated by setting the corresponding coefficients to 0s. Time and group interaction can also be added in this step.
7. Generate random error 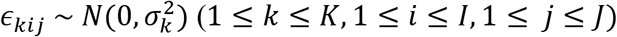, where 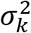 controls the within-subject correlation for taxon *k*.
8. Add covariate (*X*_*i*_), confounder (*Z*_*i*_), time (*T*_*ij*_) effects and the random effect (*ϵ*_*kij*_) using a log linear model 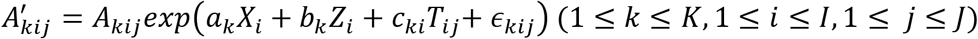.
9. Normalize into the proportion 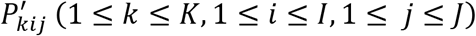 based on 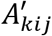. Generate the sequencing depth *D*_*ij*_ (1 ≤ *i* ≤ *I*, 1 ≤ *j* ≤ *J*) based on a negative binomial distribution. Finally, generate the read counts 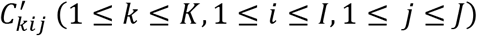 based on a multinomial distribution with parameters 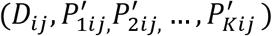.

The matched-pair and the replicate sampling design can be regarded as special cases of the longitudinal design. The matched-pair data can be generated by including two time points in the previous steps and setting the covariate effect to be 0 while the replicate sampling data can be generated by setting the time effect to be 0.

### Simulation settings for the evaluation

To comprehensively evaluate the performance of DAA-c methods for correlated microbiome data, we simulate various settings covering a wide range of signal structures (Table 1). We focus on testing the effect of *X*_*i*_ in the replicate sampling design, *T*_*i*_ in the matched-pair design, *X*_*i*_ and *T*_*i*_ in the longitudinal design. We study the performance under both the balanced and unbalanced differential settings, where the differential taxa could have random (“balanced”) or the same direction of change (“unbalanced”). The unbalanced setting creates strong compositional effects and is statistically more challenging than the balanced setting. We study the performance under both a high and low-diversity microbial community as represented by the stool and vaginal microbiome, respectively.

To further dissect the performance of the DAA-c methods, we study two levels of effect sizes under each setting, denoted as “+” and “++”, representing moderate and large effects. Since confounders are common for microbiome studies^63,64^ and adjusting confounders is critical in obtaining robust biological findings, we simulate one continuous confounder with a correlation of ∼0.6 between the covariate and the confounder. In the default setting, we include *K* = 500 taxa and a total of *n* = 200 samples. Specifically, for the replicate sampling design, we simulate 100 subjects, each with 2 replicates. For the matched-pair design, we simulate 100 subjects, each with a pre- and post-treatment sample. For the longitudinal design, we simulate 40 subjects, each with 5 time points. We also study the effect of a small sample size/taxa number by decreasing the number of subjects to 20 and the number of taxa to 50, roughly representing family- or genus-level abundance data after filtering. For all simulations, we generate sequencing depths from a negative binomial distribution with a mean depth 10,000 and a dispersion parameter 5 (*rnegbin*(theta=5) in R package “MASS”). Throughout the simulation setting, we include 10% randomly drawn differential taxa. We also let 5% and 10% taxa be affected by the confounder for the differential and non-differential taxa, respectively.

### Differential abundance analysis methods evaluated

We evaluate the widely used and recently developed DAA-c methods including ZIGMM^27^, NBMM^36^, FZINBMM^37^, GLMMadaptive^38^, glmmTMB^65^, glmer.nb^66^, GLMMPQL^66^, LDM^28,67^, LinDA^26^, ZIBR^29^ and MaAsLin2^24^. A detailed summary is shown in Table 1. For count model-based methods including NBMM, FIZNBMM, GLMMadaptive, glmmTMB, glmer.nb, and GLMMPQL, the log GMPR (geometric mean of pairwise ratios)^49^ size factors are used as the offset to account for the library size variation. We choose “family=zi.negative.binomial()” and “family=nbinom2” for GLMMadaptive and glmmTMB, respectively. For ZIGMM, a log transformation is applied before running the method and a log GMPR size factor is used as the offset. LDM currently can only be directly applied to replicate sampling and matched-pair designs, thus it is not tested for the general longitudinal design. Default settings are chosen for all the methods evaluated. For all simulated datasets, taxa with prevalence less than 10% or the maximum proportion less 0.2% are excluded from testing as is usually done in practice. For consistency, all filtering steps in the evaluated methods are disabled, and the same preprocessed datasets are used as the input to all methods.

### Performance evaluation for the simulation study

We evaluate the performance of DAA-c methods based on their ability to control for false positives and their power to detect the true associations after applying false discovery rate (FDR) control (BH procedure^68^) at the 5% target level. False positive control is assessed based on the observed empirical FDR, which is the false discovery proportion (FDP) averaged over 100 simulation runs (1,000 simulation runs for the global null). Power is assessed based on the average true positive rate (TPR). FDP and TPR are defined as:

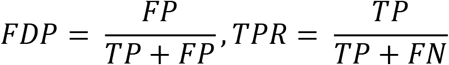

where FP, TP, and FN are the number of false positives, true positives, and false negatives, respectively. To facilitate assessment and visual interpretation, we use a scoring system to summarize the performance across settings (Supplementary Table 2):

#### False positive control scoring system

Observed FDR ∈[0,0.05], (0.05,0.1], (0.1,0.2], and (0.2,1] scores 3 green stars, 2 yellow stars, 1 red star, and 0 star (all gray), respectively. The total score is the number of stars the method receives for each setting. If an observed FDR is larger than 0.05 but its 95% confidence interval covers 0.05, we also assign 3 green stars.

#### Power scoring system

We rank the methods based on their average TPRs (higher rank, better power). The total score is the sum of the ranks for each setting.

#### Overall score

To produce an overall score, we first convert the total FDR and TPR scores into ranks (“TPR rank” and “FDR rank”). These ranks are summed for each method to produce an “overall score”. This strategy assigns equal weight to false positive control and power. The order of the methods displayed in the figures is then based on the overall score.

### Real microbiome datasets

Three real microbiome datasets representing replicate sampling, matched-pair and longitudinal desgins were used to compare the performance of competing DAA-c methods. The first dataset (“Smoker2010”) was generated to study whether smoking has an effect on the human upper respiratory tract (URT) microbiome via 16S rRNA gene-targeted sequencing^69^. Replicate sampling was used in this study (left and right nose, left and right throat). The dataset was downloaded from the Qiita database^70^ with the study ID 524. Samples with reads less than 1000 were excluded from downstream analysis. We focused on comparing the URT microbiome between smokers and non-smokers based on the two throat samples and excluded samples with less than 1,000 read counts and OTUs with a maximum proportion less than 0.002 or a prevalence less than 10% of the samples. Finally, 124 samples (31 smoking subjects and 31 non-smoking subjects, each subject has 2 replicates from the left and right side of the throat) and 197 OTUs were included in the analysis. Sex is the confounder (p=0.01) in this dataset and was included as a covariate.

The second dataset (“Nicholas2013”) was generated to study the impact of cleaning on the surface microbiome within a NICU (neonatal intensive care units)^6^. The dataset was downloaded from the Qiita database^70^ with the study ID 1798. Matched-pair design was used in this study. 16S rRNA gene-targeted sequencing was used to profile the NICU surface microbial communities before and after cleaning^6^. Genus-level abundance data were used in this analysis. Genera with a maximum proportion less than 0.002 or a prevalence less than 10% of the samples were excluded from the analysis. Finally, 70 samples (35 matched pairs before and after intensive cleaning) and 110 genera were included in the analysis.

The third dataset (“IBD2017”) was generated from a longitudinal study of the gut microbiome in Inflammatory bowel disease (IBD) patients^53^. The dataset was downloaded from the Qiita database^70^ with the study ID 1629. The fecal samples were provided by patients every third month for a two-year period. Again, 16S rRNA gene-targeted sequencing was used to profile the stool microbial community. In this analysis, we focused on comparing the gut microbiome of ICDr patients (ICD patients that had previously undergone ileocaecal resection) to healthy controls (group difference) and testing whether the longitudinal trend differed by the group (time and group interaction). Fecal calprotectin (f-calprotectin) concentration is the confounder (p<0.001) and was included as a covariate. Samples with a sequencing depth less than 10,000 and taxa with a prevalence less than 10% or a maximum proportion less than 0.002 were excluded from the analysis. Subjects with samples less than 2 were also excluded from the analysis. As a result, a total of 147 samples and 498 OTUs were included in the analysis. The 147 samples came from 9 healthy controls (2-8 longitudinal samples per subject) and 18 ICDr subjects (2-8 longitudinal samples per subject).

### Availability of data and materials

The datasets and codes supporting the conclusions of this article are available in the https://github.com/chloelulu/DAA-c repository. The semiparametric simulation approach is implemented as “*SimMSeqC*” function in the CRAN *GUniFrac* package (https://CRAN.R-project.org/package=GUniFrac). All analyses are performed in R v4.0.3 on a x86_64-pc-linux-gnu (64-bit) Red Hat Enterprise Linux Server 7.9 at Mayo Clinic.

## Acknowledgements

The work was supported by Center for Individualized Medicine at Mayo Clinic and NIH R21 HG011662, NIH R01 GM144351, NSF DMS 2113360.

## Author Contributions

J.C. conceived the idea, designed the study. L.Y. carried out the evaluation. J.C and L.Y. drafted the manuscript together.

## Competing Interests

The authors declare no competing interests.

## Materials & Correspondence

Correspondence to Jun Chen.

---

**Supplementary Table 1.** Description of the parameters used in the proposed simulation framework

| Symbol | Definition |
| --- | --- |
| $M_g^X$ | Mean of the covariate effect $a_k$ across taxa |
| $S_g^X$ | Variance of the covariate effect $a_k$ across taxa |
| $M_g^Z$ | Mean of the confounder effect $b_k$ across taxa |
| $S_g^Z$ | Variance of the confounder effect $b_k$ across taxa |
| $M_g^T$ | Mean of the subject-averaged time effect $c_k$ across taxa |
| $S_g^T$ | Variance of the subject-averaged time effect $c_k$ across taxa |
| $S_b^T$ | Variance of the subject-specific time effect $c_{ki}$ reflecting the between-subject variability |
| $\sigma_k^2$ | Variance of the random error $\epsilon_{kij}$ that controls the within-subject correlation |

**Supplementary Table 2.** Performance scoring metrics in evaluation of DAA-c methods

| False positive control |  |  | True positive rate |  |
| --- | --- | --- | --- | --- |
| FDR | Rating | Score | TPR rank | Score |
| $[0,0.05]^{\#}$ | ★ ★ ★ | 3 | Highest TPR among 11 methods | 11 |
| $(0.05,0.1]$ | ★ ★ | 2 | Second highest TPR among 11 methods | 10 |
| $(0.1,0.2]$ | ★ | 1 | ... | ... |
| $(0.2,1]$ | | 0 | Lowest TPR among 11 methods | 1 |

<sup>#</sup> Rating "\*\*\*" is given when the 95% confidence interval of the FDR estimate covers 0.05.

**Fig. S1.**
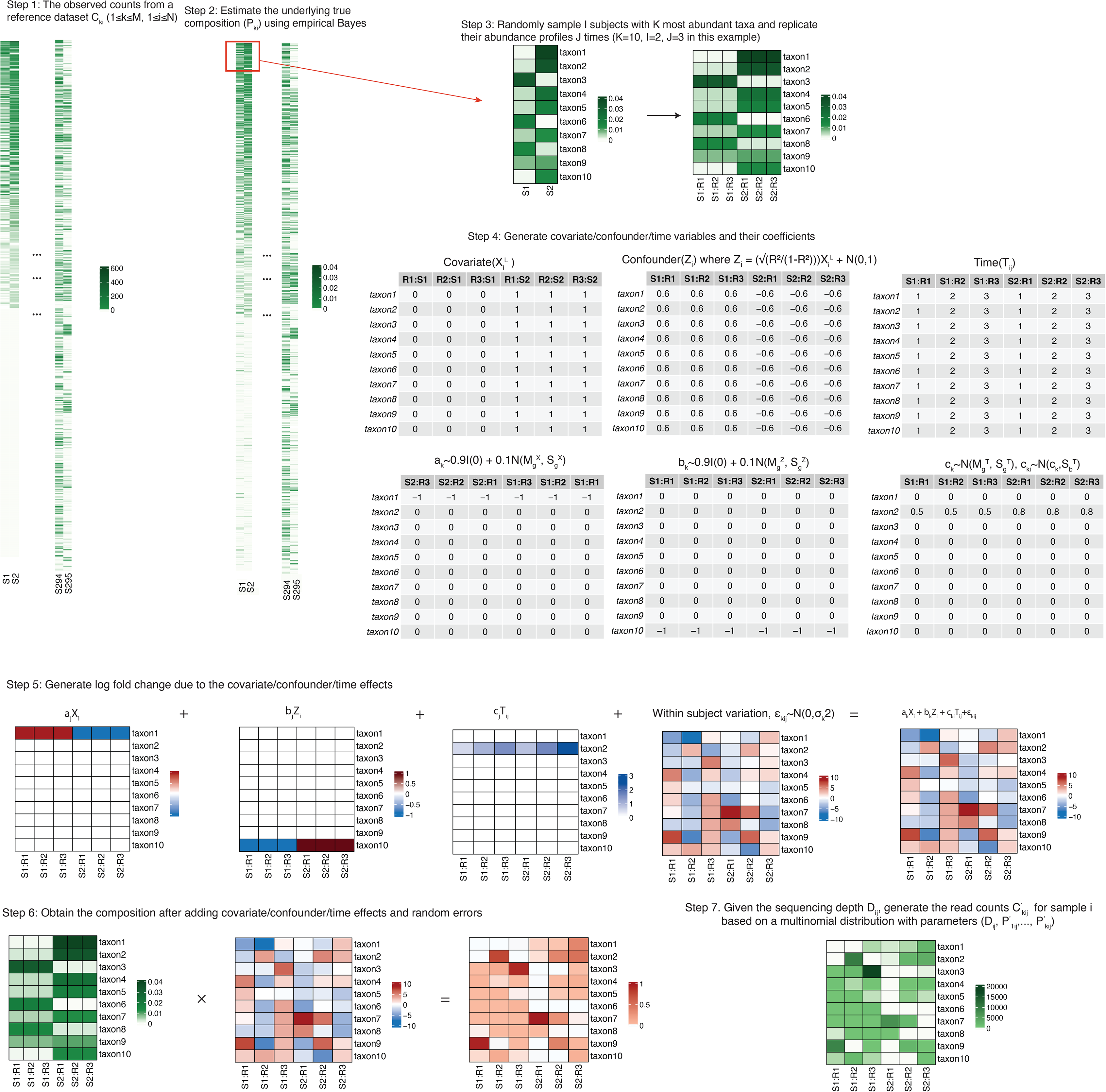
Basic steps of the proposed semiparametric simulation framework for generating longitudinal microbiome data.

**Fig. S2.**
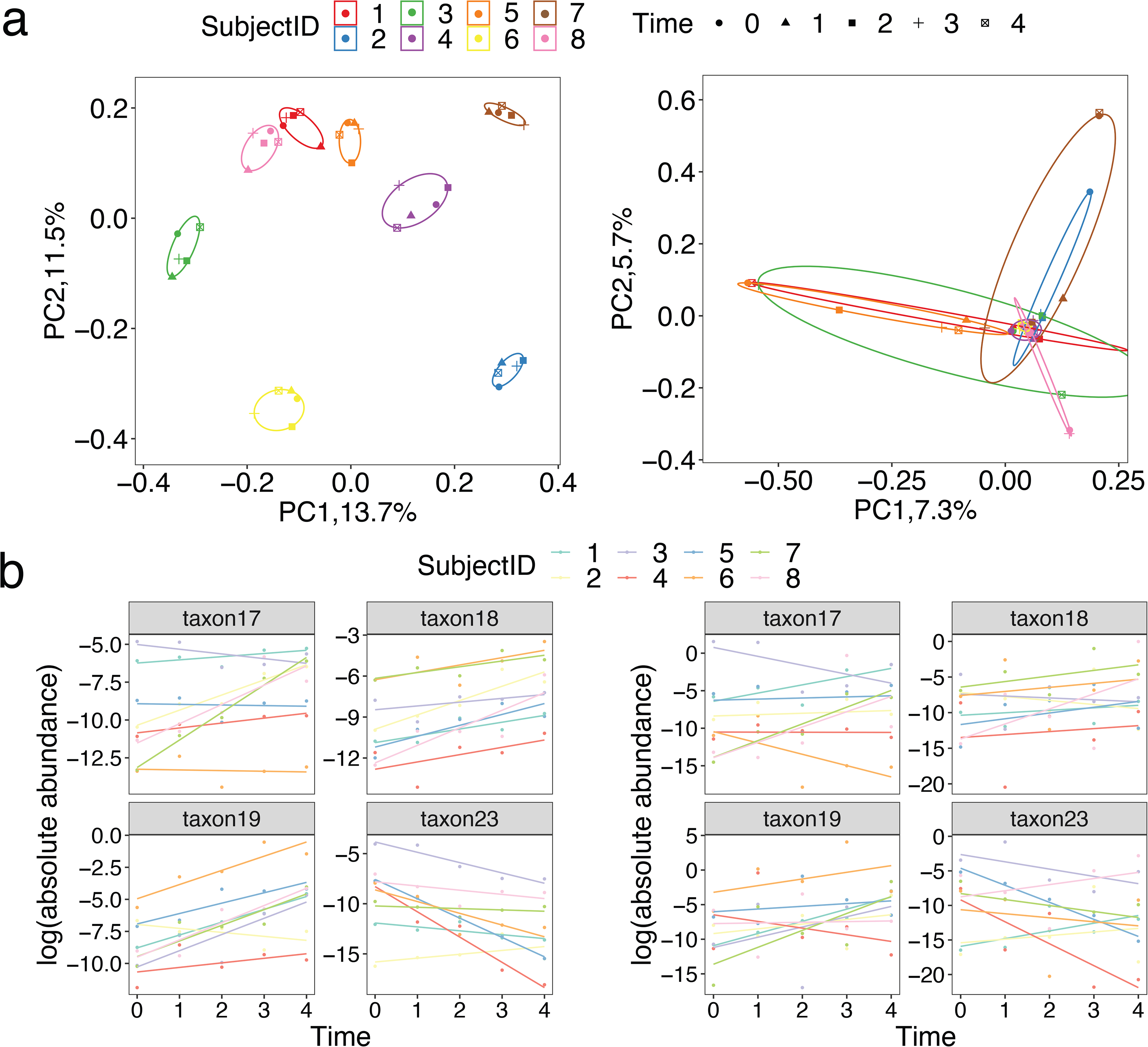
Patterns of the microbiome data generated by the proposed semiparametric framework (balanced change, stool, longitudinal sampling, I= 8, J = 5, K = 500). **a**. PCoA plot based on the Bray-Curtis distance using the data generated by the proposed simulation framework. Left: 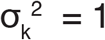, Right: 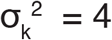. **b**. The temporal trends of the simulated abundances of 4 randomly selected taxa (random slope model). Left: 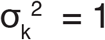, Right: 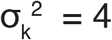.

**Fig. S3.**
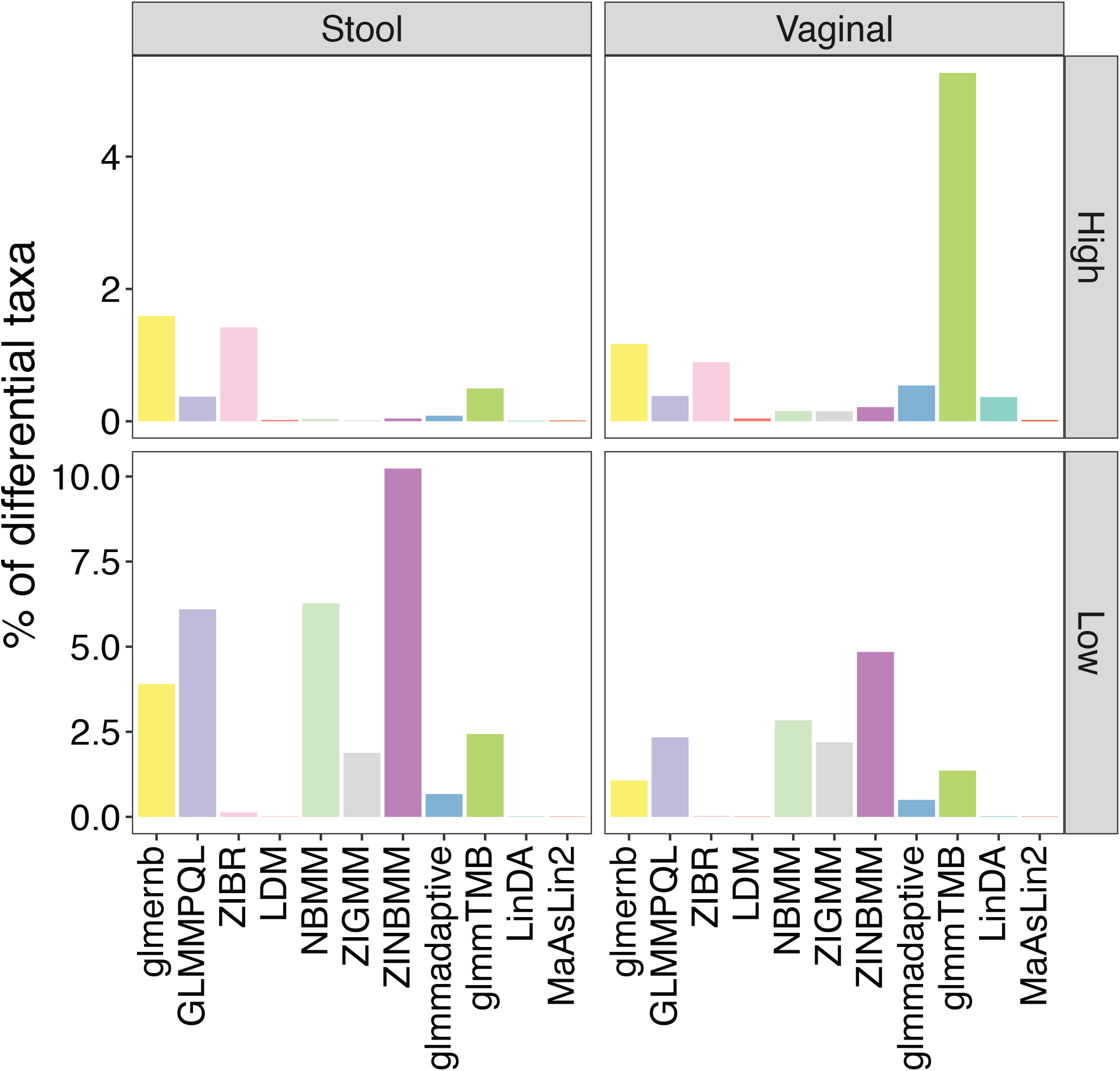
The average percentage of significant taxa at 5% FDR of DAA-c methods under the global null setting (replicate sampling).

**Fig. S4.**
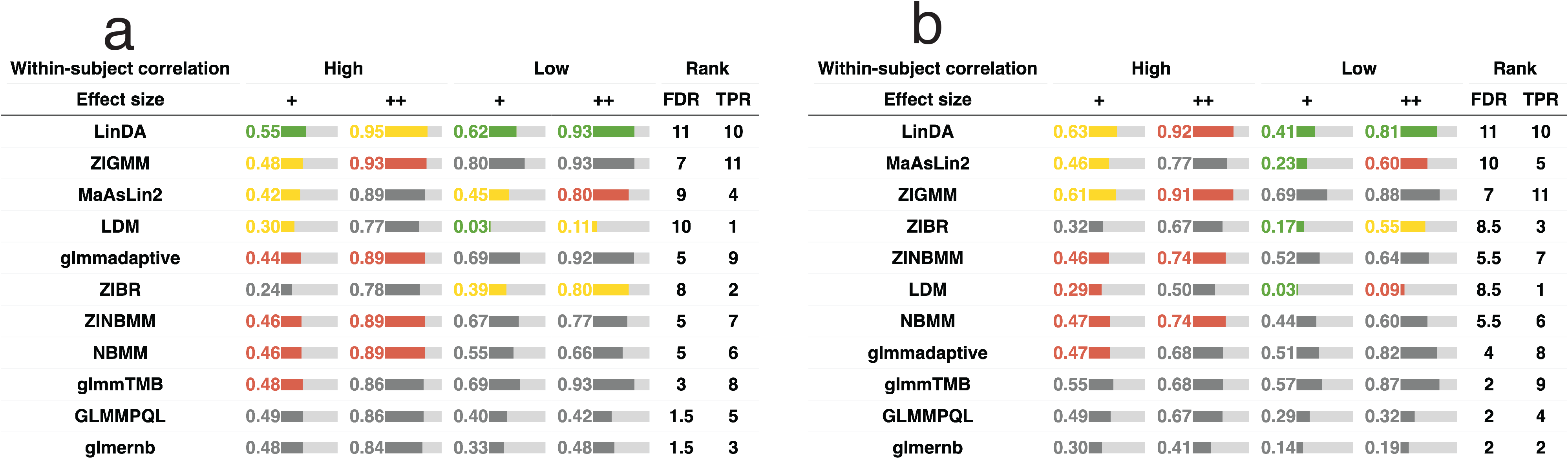
False positive control and power under the replicate sampling design (unbalanced change setting, **a**: stool and **b**: vaginal). ‘+’, and ‘++’ represent moderate and large effect sizes, respectively. “High” and “Low” within-subject correlations are simulated with 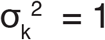 and 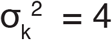, respectively. Green, yellow, red, and gray colors indicate the empirical FDR. The green color indicates that the method controls the FDR at the 5% target level (the 95% confidence interval covers 5%). Yellow, red and gray colors indicate the observed FDR level in (0.05-0.1], (0.1, 0.2], and (0.2, 1], respectively. The length of the bar is proportional to the average TPR and the actual TPR is shown in the bar. FDR and TPR ranks are based on the average FDR and TPR scores across settings. The order of the method is arranged based on the sum of the FDR and TPR ranks.

**Fig. S5.**
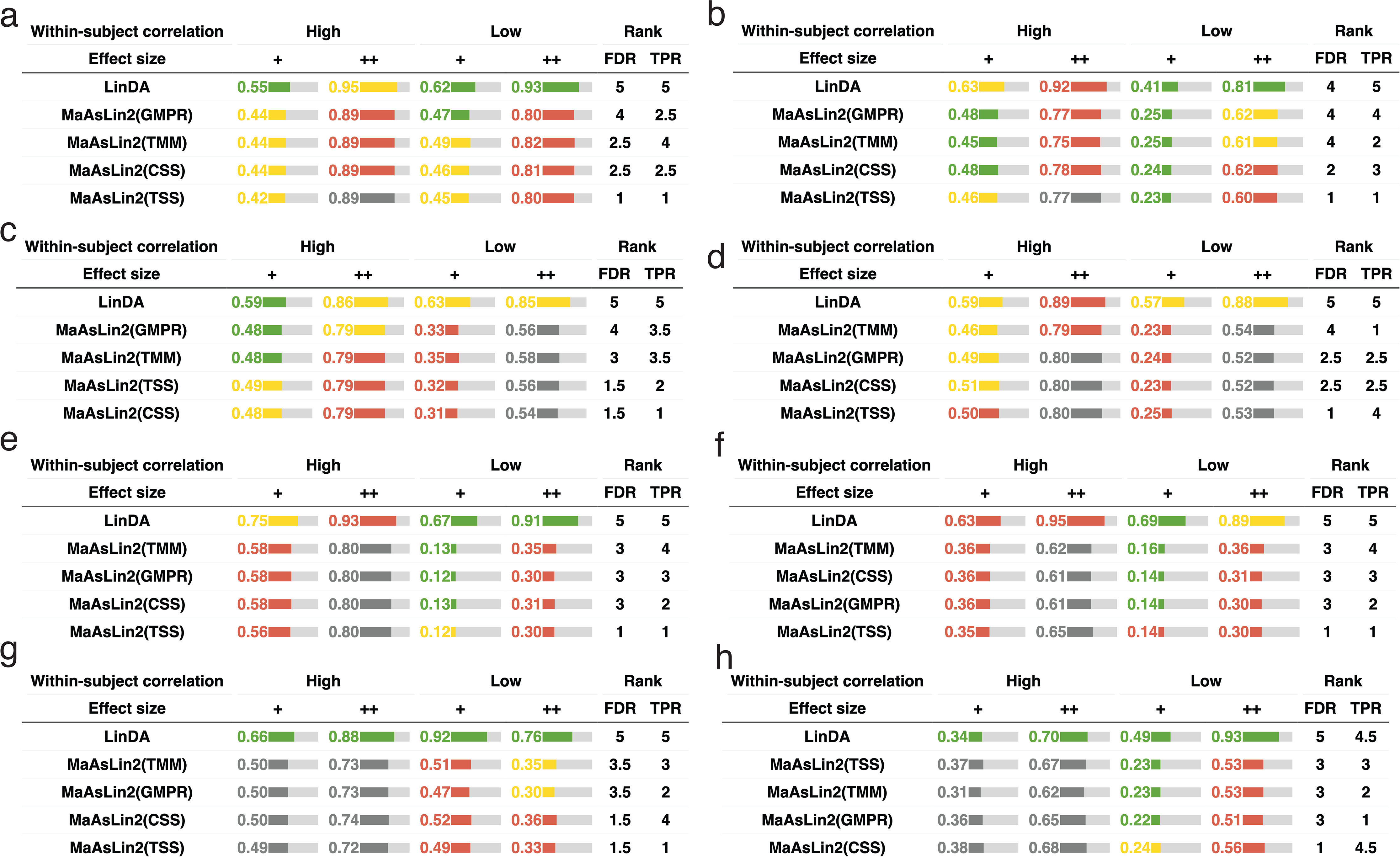
Performance comparison of MaAsLin2 with different normalization methods under unbalanced change setting (total sample size: 200, number of taxa: 500). **a,b**. False positive control and power under the replicate sampling design (**a**: stool and **b**: vaginal). **c,d**. False positive control and power under the matched-pair design (**c**: stool and **d**: vaginal). **e,f**. False positive control and power under the general longitudinal design and testing the effect of X (**e**: stool and **f**: vaginal). **g,h**. False positive control and power under the general longitudinal design and testing the effect of T (**g**: stool and **h**: vaginal). ‘+’, and ‘++’ represent moderate and large effect sizes, respectively. “High” and “Low” within-subject correlations are simulated with 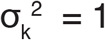 and 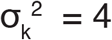, respectively. Green, yellow, red, and gray colors indicate empirical FDR. The green color indicates that the method controls the FDR at the 5% target level (the 95% confidence interval covers 5%). Yellow, red and gray colors indicate the observed FDR level in (0.05-0.1], (0.1, 0.2], and (0.2, 1], respectively. The length of the bar is proportional to the average TPR and the actual TPR is shown in the bar. FDR and TPR ranks are based on the average FDR and TPR scores across settings. The order of the method is arranged based on the sum of the FDR and TPR ranks.

**Fig. S6.**
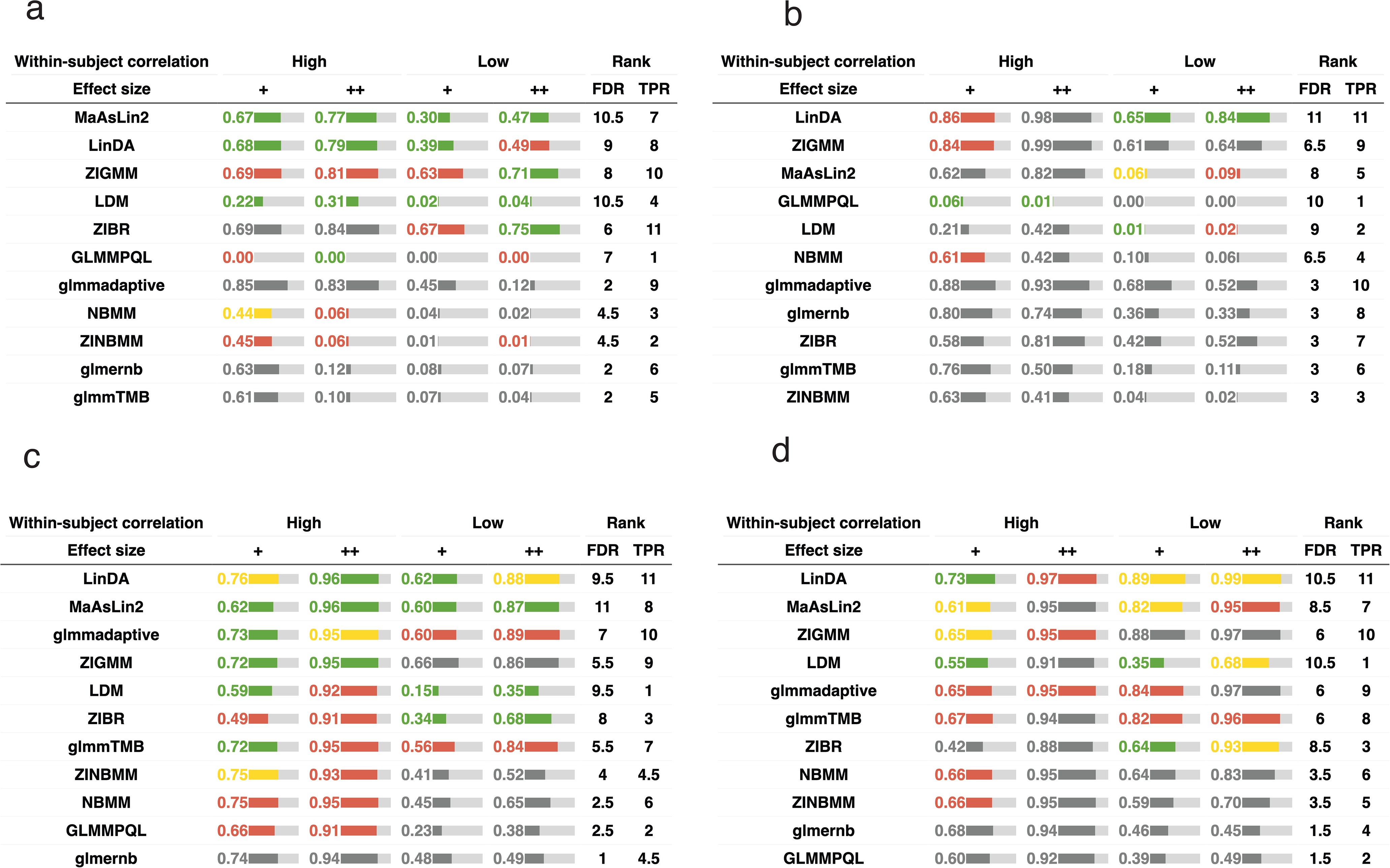
False positive control and power under a small sample size and a small number of taxa (replicate sampling, stool data). **a,b**. False positive control and power under a small sample size under (40 samples, **a**: balanced change setting and **b**: unbalanced change setting). **c,d**. False positive control and power of a small taxa number (50 taxa, **a**: balanced change setting and **b**: unbalanced change setting). ‘+’, and ‘++’ represent moderate and large effect sizes, respectively. “High” and “Low” within-subject correlations are simulated with 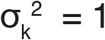 and 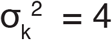, respectively. Green, yellow, red, and gray colors indicate empirical FDR. The green color indicates that the method controls the FDR at the 5% target level (the 95% confidence interval covers 5%). Yellow, red and gray colors indicate the observed FDR level in (0.05-0.1], (0.1, 0.2], and (0.2, 1], respectively. The length of the bar is proportional to the average TPR and the actual TPR is shown in the bar. FDR and TPR ranks are based on the average FDR and TPR scores across settings. The order of the method is arranged based on the sum of the FDR and TPR ranks.

**Fig. S7.**
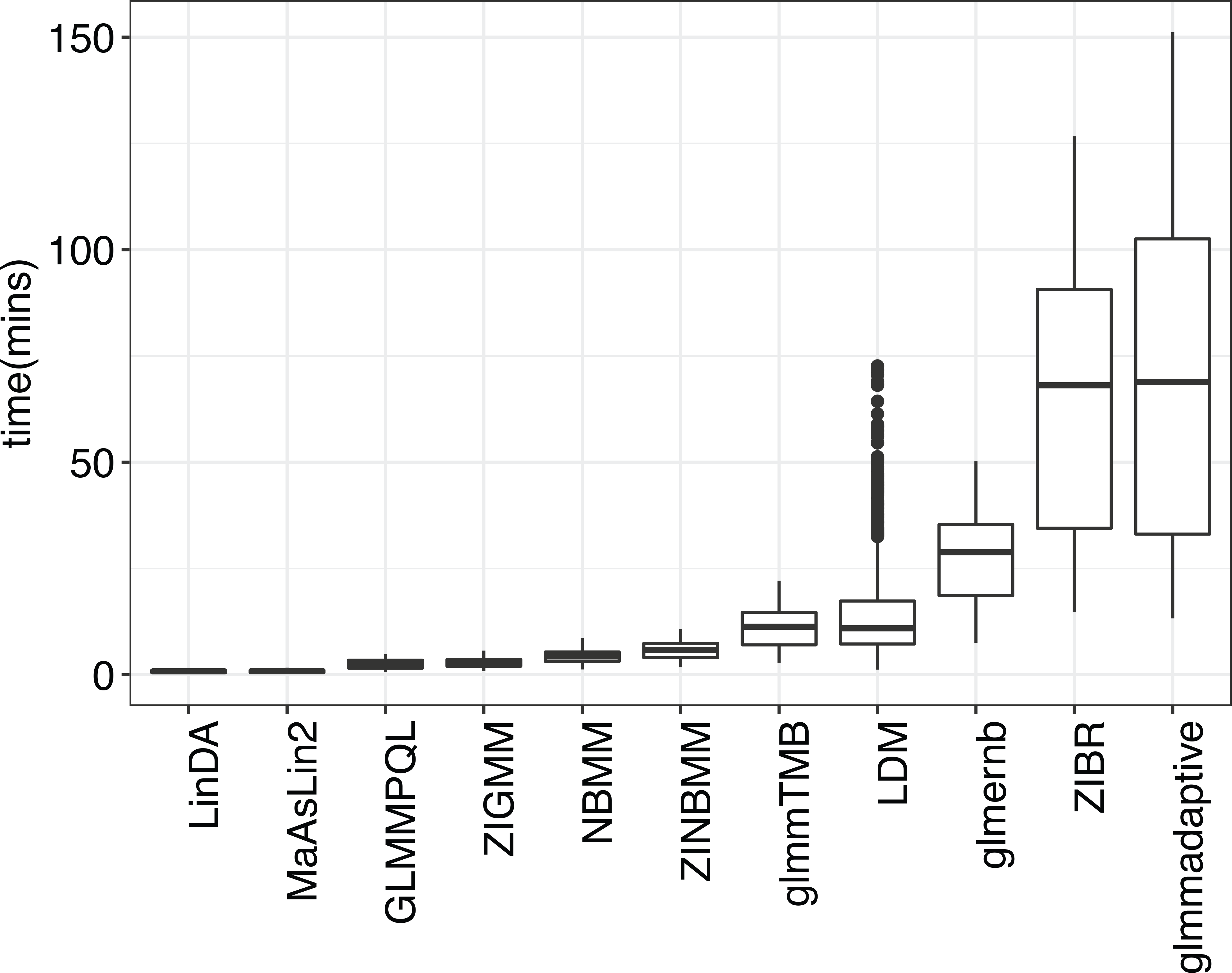
Run times (x86_64-pc-linux-gnu (64-bit) Red Hat Enterprise Linux Server 7.9, Intel(R) Xeon(R) CPU E5-2698 v4 @ 2.20GHz, 8GB running memory) of the evaluated DAA-c methods over simulation runs (balanced and unbalanced setting, stool and vaginal data, 200 samples and 500 taxa).

**Fig. S8.**
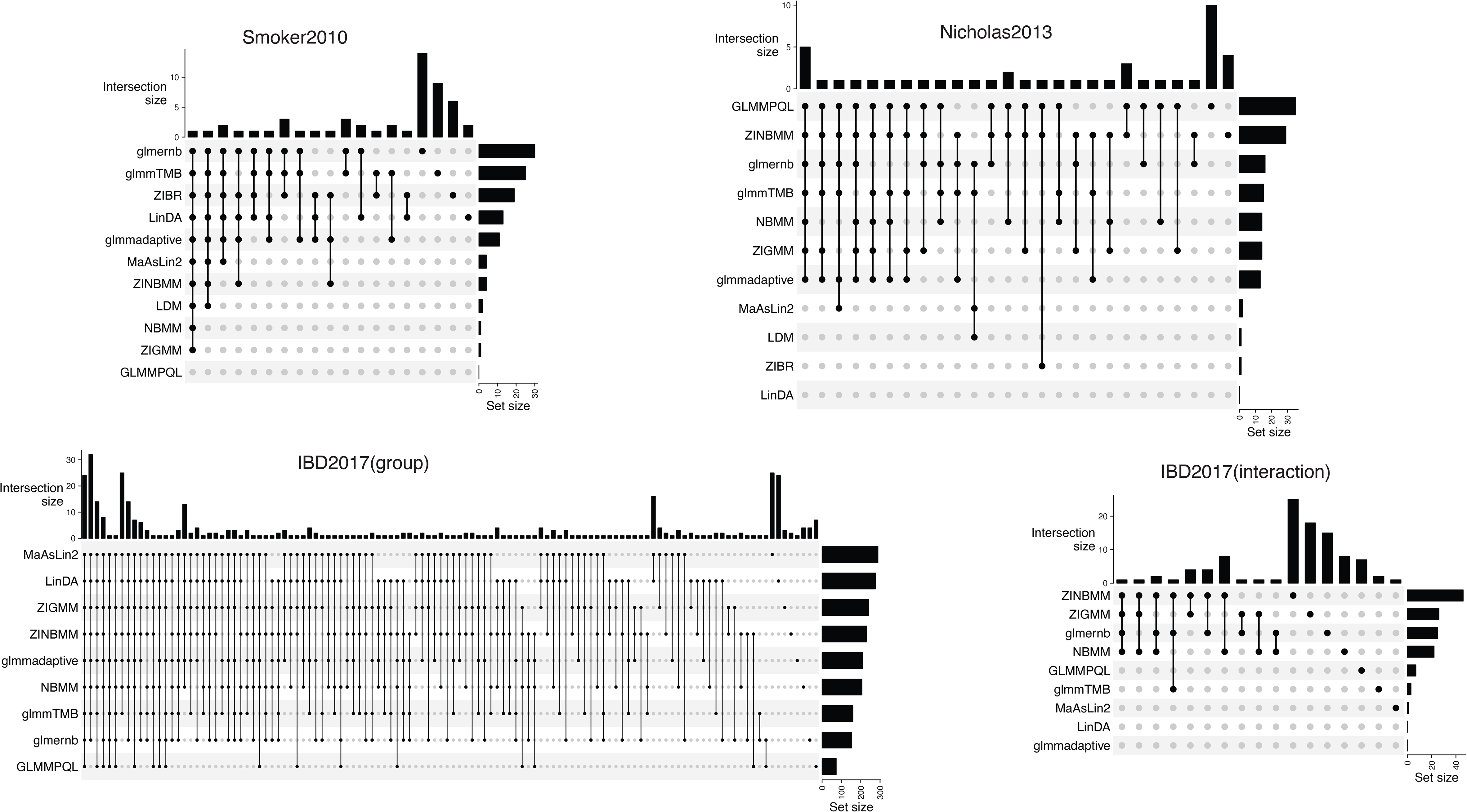
Overlaps of significant taxa between DAA-c methods on the real datasets at 10% FDR. Set size means the total number of differential taxa discovered by each method. Intersection size means the number of differential taxa commonly found by the methods indicated by the dots.

## Notes

### Competing Interest Statement

The authors have declared no competing interest.

